# Nucleus accumbens neurons encode initiation and vigor of reward approach behavior

**DOI:** 10.1101/2021.01.12.425739

**Authors:** David Levcik, Adam H. Sugi, José A. Pochapski, Gabriel Baltazar, Laura N. Pulido, Cyrus Villas-Boas, Marcelo Aguilar-Rivera, Romulo Fuentes-Flores, Saleem M. Nicola, Claudio Da Cunha

## Abstract

The nucleus accumbens (NAc) is considered an interface between motivation and action, with NAc neurons playing an important role in promoting reward approach. However, the encoding by NAc neurons that contribute to this role remains unknown. Here, we trained male rats to find rewards in an 8-arm radial maze. The activity of 62 neurons, mostly in the shell of the NAc, were recorded while rats ran towards each reward place. General linear model (GLM) analysis showed that variables related to the vigor of the locomotor approach, like speed and acceleration, and the fraction of the approach run completed were the best predictors of the firing rate for most NAc neurons. Nearly 23% of the recorded neurons, here named locomotion-off cells, were inhibited during the entire approach run, suggesting that reduction in firing of these neurons promotes initiation of locomotor approach. Another 24% of the neurons presented a peak of activity during acceleration followed by a valley during deceleration (peak-valley cells). Together, these neurons accounted for most of the speed and acceleration encoding identified in the GLM analysis. Cross-correlations between firing and speed indicated that the spikes of peak-valley cells were followed by increases in speed, suggesting that the activity of these neurons drives acceleration. In contrast, a further 19% of neurons presented a valley during acceleration followed by a peak just prior to or after reaching reward (valley-peak cells). These findings suggest that these three classes of NAc neurons control the initiation and vigor of the locomotor approach to reward.

**Significance Statement:** Deciphering the mechanisms by which the NAc controls the vigor of motivated behavior is critical to better understand and treat psychiatric conditions in which motivation is dysregulated. Manipulations of the NAc profoundly impair subjects’ ability to spontaneously approach reward-associated locations, preventing them from exerting effort to obtain reward. Here, we identify for the first time specific activity of NAc neurons in relation to spontaneous approach behavior. We discover three classes of neurons that could control initiation of movement and the speed vs. time trajectory during locomotor approach. These results suggest a prominent but heretofore unknown role for the NAc in regulating the kinematics of reward approach locomotion.

## Introduction

Since the early anatomical and physiological studies that established the nucleus accumbens (NAc) as a limbic-motor interface (Mogenson et al., 1980), a consensus view has emerged that a major function of NAc neurons is to promote the vigorous pursuit of rewards (Nicola, 2007; Salamone and Correa, 2012; Nicola, 2016). Manipulations of the NAc, and particularly of its dopamine input, impair performance of high-effort operant tasks while leaving lower-effort tasks relatively unaffected (Salamone et al., 1999), bias animals to choose the less effortful option in T-maze tasks (Salamone et al., 1994; Cousins et al., 1996; Hauber and Sommer, 2009), and reduce the probability of engaging in locomotor approach responses to reward-predictive cues (Nicola, 2010; Ambroggi et al., 2011). Consistent with these observations, many NAc neurons are excited by reward-predictive cues (Nicola et al., 2004; Gmaz et al., 2018), and these excitations predict the vigor of the approach response – specifically, the firing is greater when the latency to initiate approach will be shorter and the speed of approach greater (McGinty et al., 2013; Morrison et al., 2017). This form of encoding likely causally contributes to vigorous performance of cued approach tasks (du Hoffmann and Nicola, 2014; Caref and Nicola, 2018).

Although NAc cue-evoked firing responses compellingly link reward prediction to effort exertion, most paradigms that have revealed an effect of NAc manipulations on effort-based performance have not involved presentation of explicit predictive cues (Cousins et al., 1996; Aberman and Salamone, 1999; Salamone et al., 1999; Hauber and Sommer, 2009). For example, NAc disruption selectively impairs high-effort operant task performance by preventing the subject from approaching the operandum after pauses in performance, which are more frequent in higher-effort than lower-effort operant tasks (Nicola, 2010). These observations suggest that NAc neurons control spontaneous approaches to rewarded locations. The NAc neuronal activity underlying this form of effort exertion remains poorly understood. One challenge is to identify the onset of spontaneous approach events, which is difficult when standard operant chambers are used because short-distance spontaneous approaches are similar to the frequent non-approach movements exhibited by rodents when they are not engaged in the task. In contrast, approach movements are readily identifiable in maze or runway tasks in which subjects must move long distances (e.g. 1 m or greater) to rewarded locations. Although NAc neuronal activity has been measured in such tasks (Shibata et al., 2001; Mulder et al., 2004; Mulder et al., 2005; German and Fields, 2007; Khamassi et al., 2008; van der Meer and Redish, 2009; van der Meer et al., 2010), the relationship between neuronal activity and locomotor vigor has not been systematically investigated, despite anecdotal reports that the speed of locomotion is reported by NAc neurons (Sjulson et al., 2018).

To clarify how NAc neurons represent the vigor of spontaneous locomotion, we recorded from NAc neurons as rats performed an 8-arm radial maze task in which 3 arms were consistently rewarded with either the same or different chocolate milk reward values. We found that speed and proximity to the rewarded target were prominently encoded by many NAc neurons during reward approach, and that neuronal activity predicted speed approximately 100 ms in advance. In addition we found 3 classes of neurons with different patterns of activity that could control when the approach run starts as well as the timing of acceleration and deceleration.

## Materials and Methods

### Subjects

We used 5 adult male Wistar rats that were three months old at the beginning of the experiment. The rats came from the breeding colony of the Federal University of Parana State and were housed in groups of 4 per cage during behavioral training and individually after surgery. The rats were maintained in a temperature-controlled room (22 ± 2 °C) with a 12-hr light/dark cycle (lights on at 7:00 am). Access to water was allowed for one hour per day, and access to food was restricted to maintain the rats’ body weight at 90% of their free-feeding weight (290 - 340 g). All experimental procedures were in agreement with the Brazilian and International legislation for animal care (Law N° 11.794 of October 8, 2008; EC Council Directive of November 24, 1986; 86/609/EEC). The project was approved by the Animal Care and Use Committee of the Federal University of Parana State and efforts were made to minimize the number of animals used, and their suffering and discomfort during the experimental procedures.

### Apparatus

We used a stainless steel eight-arm radial maze, which had a surface covered with black contact paper, for behavioral training. Each arm (62 cm x 13 cm) and the central platform (an octagon with a 30 cm diameter) were elevated 60 cm above the floor. The walls of all arms were 5 cm high. One or four drops (25 µl per drop) of chocolate milk were delivered at the ends of reward arms before the start of each trial. A white curtain was installed around the maze and several salient visual cues (black felt geometrical shapes) were attached to it and remained in the same locations throughout behavioral training and experiments. Four light bulbs (15 W) were spaced equally on a metal frame (70 × 70 cm) above the center of the maze. This metal frame also served as a support for a motorized commutator (Plexon, USA), a camera (Allied Vision Technologies GmbH, Germany) connected to the main computer, and an amplifier (Plexon, USA). An OmniPlex D Neural Data Acquisition system (Plexon, USA) was located in the same room. Metal mesh was installed on the walls of the experimental room and grounded to create a Faraday cage.

### Behavioral procedures

All rats were handled (5 min/day) by the experimenter for three consecutive days before the start of behavioral pre-training. During pre-training, rats were habituated to the maze and the chocolate milk reward by placing them at the end of one arm and letting them drink the chocolate milk from the reward receptacle.

After the pre-training phase, rats were trained to collect drops of chocolate milk consistently located at the ends of the same three arms of an eight-arm radial arm maze (Fig. 1A). The positions of these three reward arms were counterbalanced among rats and were chosen in a pseudorandom manner so that two adjacent arms were never baited. In the same-reward group (n = 2), all reward arms (X, Y and Z) contained 4 drops of chocolate milk (100 µl) in 100% of trials. In the different-reward group (n = 3), one of the reward arms contained four drops (100 µl) of chocolate milk in 100% of trials (high reward, H arm), another reward arm contained four drops (100 µl) in 66.7% of trials (medium reward, M arm), and the last reward arm contained one drop (25 µl) in 100% of trials (low reward, L arm). The arm that was less preferred by the rats at the end of the pre-training phase was assigned as the H arm, and the most preferred arm was assigned as the L arm. The rewarded locations, and the reward amounts and probabilities, remained constant for a given rat throughout training and experiments.

**Figure 1:**
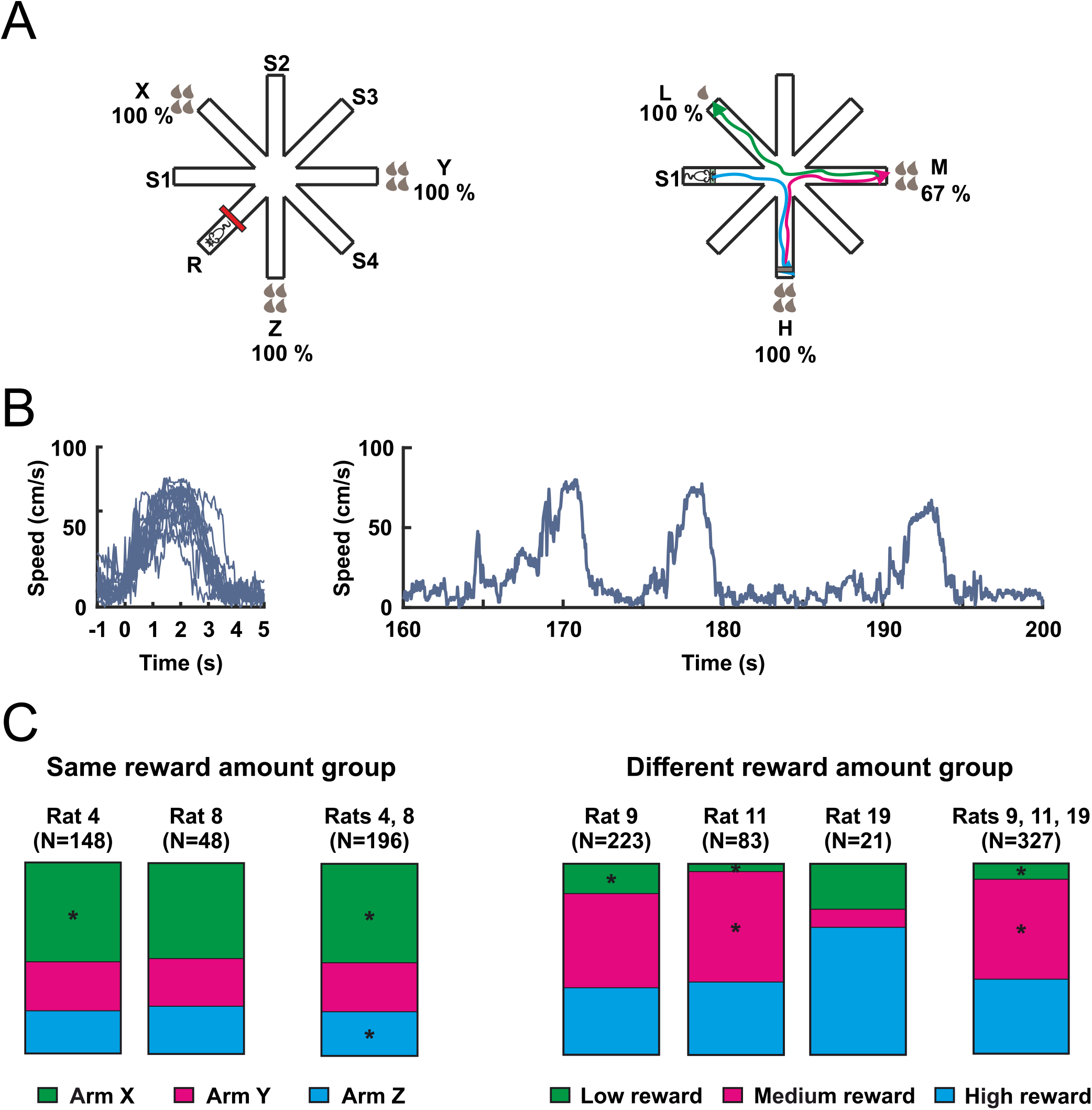
The eight-arm radial maze task. **(A)** Rats were trained to find drops of chocolate milk in the ends of the 3 baited arms. Two rats were trained to find the same reward amount (4 drops) with 100% probability (left) and the other three rats a different reward amount with a different probability (middle) in the end of the three reward arms. Each brown drop represents 25 μl of chocolate milk. The probabilities of reward occurrence are displayed next to each reward arm. Arms X, Y, and Z had equal reward value, and arms L, M, and H had low, medium, and high reward values, respectively. The red line represents an obstacle that was placed in the middle of the resting arm between trials to keep the rat restricted in the distal half of the starting arm. S1, S2, S3 and S4 indicate the starting arms. **(B)** Superimposed changes in speed during all runs of an individual session (left). Changes in speed during three subsequent runs (right). **(C)** Bars show the fraction of trials in which subjects entered the indicated arm first. * P < 0.01, compared to random choice (Fisher test).

In both same-reward and different-reward groups, a fourth arm was consistently used as a resting platform, where the rats were placed and restricted between trials. The other four arms were used as starting positions (S1 - S4, Fig. 1A). The rats underwent nine trials per day. Two starting positions were alternated in a pseudorandom order, one used three times and the other two times. Each trial finished when all rewards were collected or after 5 min elapsed. The rats were trained five days per week until they reached the following criteria in three consecutive training days: a) no more than 20% reference memory errors (entering a non-reward arm); b) the high reward arm (4 drops, 100% probability) is the last choice in no more than 20% of trials; and c) the low reward arm (1 drop, 100% probability) is the first choice in no more than 20% of trials. Criteria b and c were applied only to the different-reward group. The rats took 40 to 50 training days to achieve these criteria.

Afterwards, rats underwent surgery to implant recording electrodes. After recovery, they were re-trained to the pre-surgery level of performance, which took approximately seven days. During these re-training sessions and the following test sessions, 12 trials per session were carried out.

### Arm preference analysis

To show potential preferences for particular reward arms among individual rats, we compared the total number of first reward arm choices from all recording sessions to the chance level (total number of trials/3 – because we used three reward arms; Fisher’s exact test).

### Surgery

The rats were anesthetized with isoflurane (2%), placed in a stereotaxic frame (David Kopf, USA), and a customized microdrive with 2 × 4 stereotrode arrays (17 μm Ni-Chrome wire with insulation; A-M Systems, USA) was chronically implanted unilaterally above NAc shell at the following coordinates: AP +1.2 to +2.2 mm from bregma, ML +0.6 to +1.0 mm from midline, DV −5.0 mm from dura. The coordinates were determined according to the Paxinos and Watson Atlas (2007). Six anchoring stainless steel screws were placed in the skull to attach the implant using acrylic dental cement. One of the screws also served as a ground. After surgery, enrofloxacin (20 mg/kg; i.m.) was injected, neomycin gel was applied to the tissue around the implant, and ibuprofen (50 mg/500 ml) was added to the drinking water for the three following days. The rats were allowed to recover for seven days after the surgical procedures.

### Electrophysiological recording

After one week of recovery, the stereotrodes were lowered in 40 μm increments (up to 160 μm/day) until single-unit activity was reliably detected and isolated. All stereotrodes were previously gold-plated with a nanoZ impedance tester (Multi Channel Systems, USA) to impedances between 100 − 300 kOhms. Two light-emitting diodes attached to the 16-channel headstage (20 x gain) signaled the position of the animal and allowed video tracking (30 frames/s) with the Cineplex system (Plexon, USA). Neural activity was amplified (1000 x), filtered (300 − 6000 Hz) and sampled at 40 kHz with the OmniPlex D Neural Data Acquisition system (Plexon, USA) to acquire and record single unit activity.

### Spike sorting

Recorded spikes were manually sorted offline into clear clusters of putative cells with Offline Sorter software (Plexon, USA). To be accepted for spike sorting, spike amplitudes had to be at least three standard deviations higher than the background activity. Clusters were based on waveform properties identified through principal component analysis. Inter-spike interval histograms were calculated, and those clusters with similar waveform shape that showed a clearly recognizable refractory period (> 1.5 ms) were considered as originating from a single neuron.

### Neuronal activity analysis

A test session consisted of 12 trials. In each trial, the rats should visit the three reward arms. The rats usually approached the reward areas of the three reward arms only once. Each approach was defined as a run – when the rat ran from the distal end of one arm towards the reward area of a reward arm (Fig. 1A). Trained rats performed few visits to non-reward arms, and only data from approaches to reward arms were analyzed. For most analyses, the time window analyzed for each run extended from 1 s before locomotion onset to 1 s after the locomotion end. Taking advantage of the fact that the animals’ behavior across runs was generally consistent, locomotion onset was defined as the first time point in which the speed of the animal reached 20 cm/s before the peak speed of the run (Fig. 1B). Locomotion end was set as the first point at which the speed dropped below 10 cm/s. We restricted our analysis to stereotyped trajectories to the reward defined by an efficiency criterion, which was calculated as the ratio of the distance traveled in the run to the length of the ideal path (from the end of one arm to the reward area of another arm). Incomplete runs, i.e. the rat enters the arm but does not reach the end of it, result in ratios <1. To eliminate most such runs, and runs in which the animal deviated substantially from a direct trajectory to the end of the reward arm, runs with high efficiency (ratio < 0.8) or low efficiency (ratio > 1.2) were discarded.

### General linear model (GLM)

The goal of the multiple regression analysis was to evaluate which performance-related parameters can be used to predict neuronal firing rate. A general linear model (GLM) was calculated with the firing rate (spikes/100-ms bin) as the dependent variable and 12 parameters as predictors (independent variables; Tab. 1). Pearson correlations (r) among pairs of predictor variables were calculated and, for the pairs that showed *r* > 0.8, one of them was excluded (e.g. time and fraction of the run; Fig. 2).

**Figure 2:**
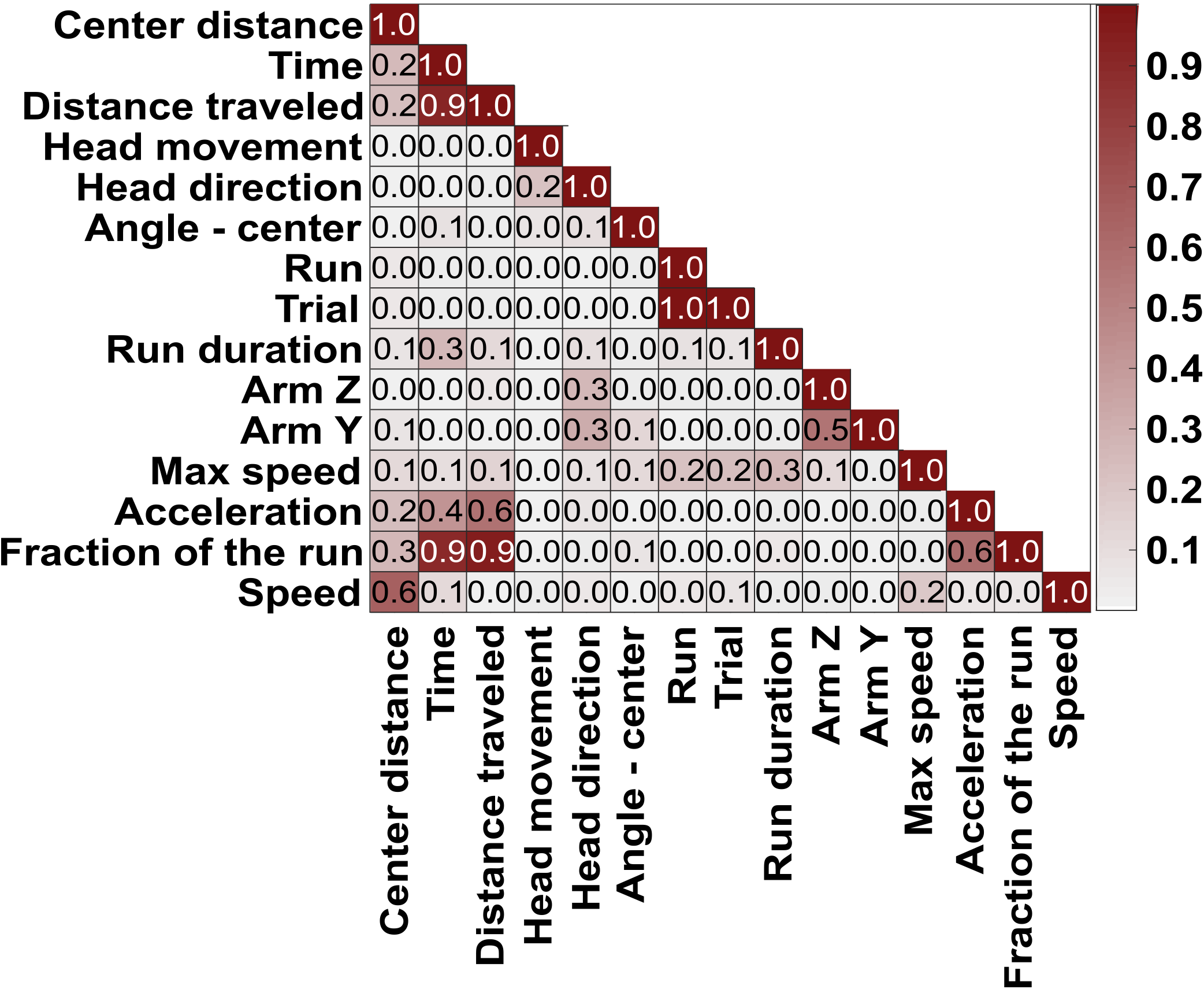
Correlation table for predictor variables considered for use in the GLM encoding model. For the pairs with high correlation (r > 0.8), one of the pairs was chosen to be included in the model.

Different continuous variables were extracted from the coordinates tracked in the recorded videos. The rat’s locations were tracked at 30 frames per second. However, to match the neuronal data that were binned at 100 ms, an interval of 3 frames was used (3 frames = 100 ms interval). Firing rate, speed and acceleration were smoothed by a sliding average window of 300 ms (Kropft et al., 2015; Rueda-Orozco and Robbe 2015). The β values were calculated with the MATLAB *glmfit* function. The multiple regression assumed a normal distribution for the data and the identity link function was used to estimate the β value for each parameter.

The GLM followed the equation:

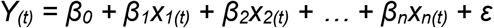

where *Y*_*(t)*_ is the predicted firing rate at time *t, β*_*n*_ is the weight of the predictor *n, x*_*n(t)*_ is the value of the predictor *n* at time *t*, and *ε* is the error term. For each neuron, the model was trained and tested by the fivefold cross validation method (Engelhard et al., 2019) as follows. The data from all the runs (speed, acceleration, firing rate, etc.) were divided into 5 parts with the same number of runs. The runs used in each part were randomly selected. The model was trained and tested five times with 4/5 of the total data, and the ability of the model to predict the remaining 1/5 of the data was assessed by correlating the predicted data to the actual data. Always, a different combination of the data was used as training and test data (Fig. 3).

**Figure 3:**
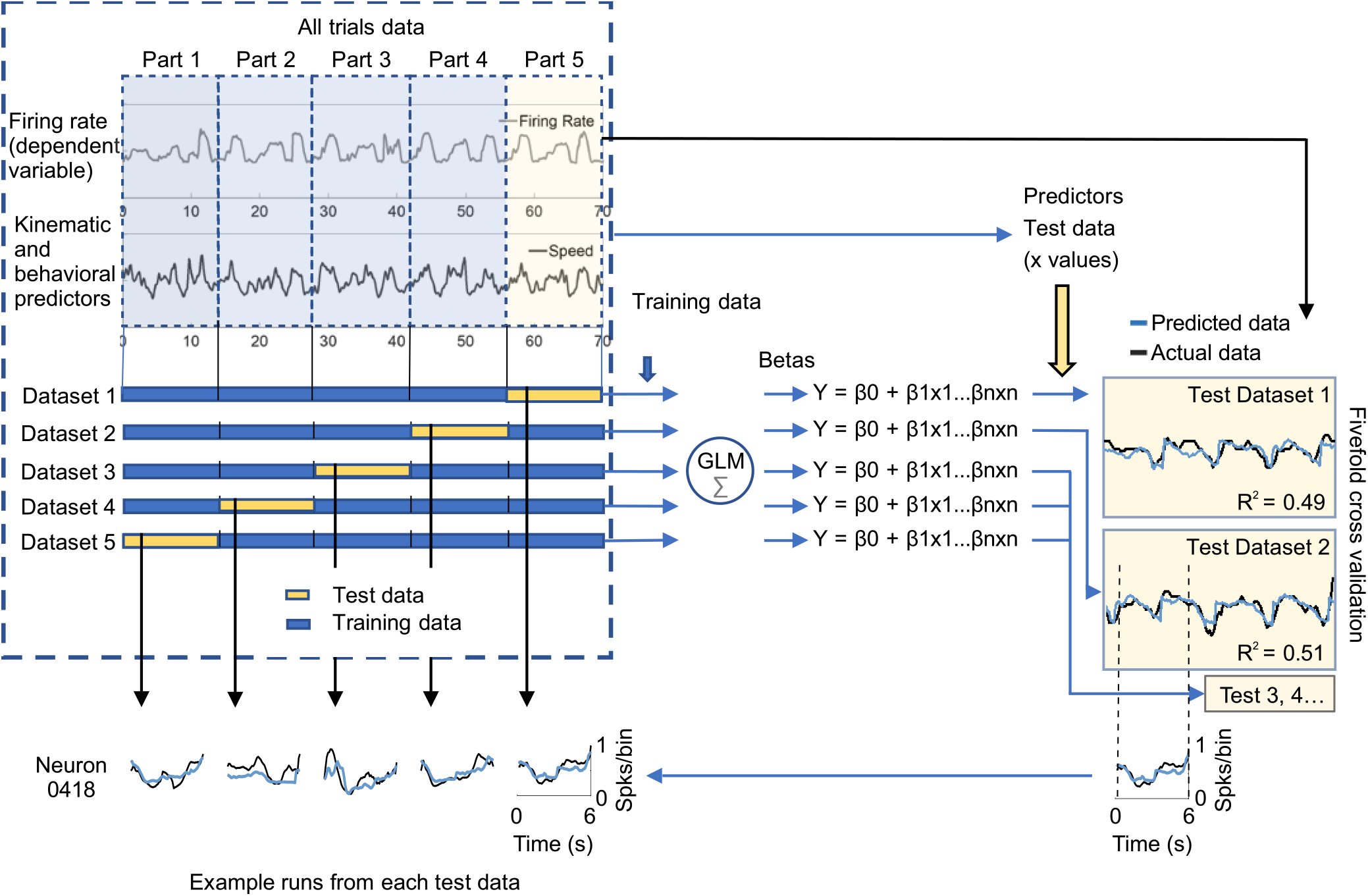
Fivefold cross validation method for evaluating the GLM encoding model. Data (firing rate and predictor variables) from a whole session were divided into five parts (datasets). Each dataset had the same number of runs (selected randomly). Four of the five datasets were used as training datasets to calculate the *β* values for each predictor. Then, using the predictors data from the remaining dataset (the test dataset) as input to the encoding model, predicted firing rates (Y) were calculated. The predicted firing rates were compared to the actual firing rates from the test data set. The variance explained by the GLM was calculated as the r^2^ of the correlation between the predicted and actual data. This process was repeated 5 times, with each dataset used only once as test data. The variance explained by the model was the mean of the 5 r^2^ values (r^2^_mean_).

The mean of the r^2^ values obtained from the five correlations (r^2^_mean_) was considered the mean fraction of variance explained by the full model (FM, the model with all predictors included). The significance of the model was evaluated by comparing the r^2^_mean_ to the distribution of r^2^_mean_ values generated by models trained with shuffled neuronal data (100 shuffled models per neuron, n = 62 neurons). Neurons for which the model explained the actual data better than the 99^th^ percentile of the shuffled models were classified as having their firing rates significantly explained by the encoding model.

### Contribution of each predictor to the GLM

To evaluate which variables had a significant contribution to the full model (FM), only neurons significantly explained by the FM were analyzed. To calculate the contribution of each of the variables (var) to the FM, the FM was compared to the full model less the selected variable (FM – var). The fraction of the variance of the FM explained by each variable was calculated as:

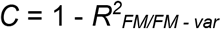

in which *C* is the fraction of the contribution of the variable to the FM and *R*^*2*^_*FM/FM – var*_ is the r^2^ of the correlation between the output of the FM and the output of the FM – var. Next, we tested whether the percent of the variance explained after each variable was removed (C*100) was significantly different from the percent of variance explained by the FM. Log(C*100), but not C*100, passed the D’Agostino & Person test of normality (D’Agostino, 1986). Therefore, log(C*100) was used for t-tests in this analysis. The contribution of a variable was considered significant if P < 0.01.

### Cross-correlogram analysis

The aim of this analysis was to test whether changes in firing rate preceded or followed changes in the animal’s speed. First, based on each neuron’s Pearson’s correlation between speed and firing rate, we divided neurons into two groups, those with positive correlations between firing rate and speed, and those with negative correlations. Only neurons with correlations higher or lower than the 99.5th percentile of a bootstrapping distribution of Pearson’s correlations were included in these groups (1000 correlations of shuffled spike times and speed were generated for each neuron). For each of the included neurons, the average speed before and after each spike (bins of 33 ms) was calculated to generate a normalized (z-scored) cross-correlogram. Finally, the time of the peak of each cross-correlogram was identified and then compared to zero (one-sample t-test).

### The two Gaussians model (2G)

The 2G model used fraction of time as the only independent variable. Time during the run was normalized by dividing the periods from locomotion onset to the peak of speed and from the peak of speed to locomotion end into 10 bins each. On average, the bin width was ∼150 ms. The time windows starting 1.05 s before locomotion onset to locomotion onset, and from locomotion end to the 1.05 s after locomotion end, were each divided in 7 bins (150 ms each). The firing rate during each bin was calculated and z-transformed.

The 2G model followed the equation:

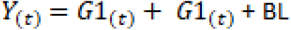

where *Y*_*(t)*_ is the predicted firing rate at time *t*, G1 is the Gaussian equation for the peak of the curve, G2 is the equation for the valley of the curve, and BL is the baseline activity.

The Gaussian equations (G_(t)_) used were:

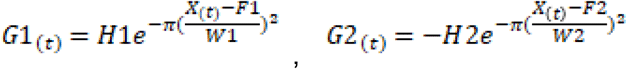

where H, F, and W are constants with the following meaning: H represents the amplitude of the Gaussian curve, which reflects the height (H1) and depth (H2) of the activity peak and the valley, respectively. F is the bin where the Gaussian is centered. Therefore, F1 is the bin in which the neuron is most active, and F2 is the bin in which the neuron is least active. W is the standard deviation of each Gaussian, which reflects the durations of the activation peak (W1) and valley (W2). For each neuron, data from all runs were randomly split in two pools. One data pool was used to adjust these constants and the other pool was used to test the prediction power of the model. 2G constants were adjusted by using the Generalized Reduced Gradient (GRG) engine solver of Microsoft Excel to minimize the sum of the square roots of the difference between the firing rate recorded in each fraction of time and the firing rate predicted by the 2G model. Next, the r and P for the correlation between the firing rate predicted by the 2G model and the firing rate activity calculated with the other data pool was calculated. The 2G model was considered to significantly predict activity of a neuron when P < 0.01.

### Data and code accessibility

The spike-train data, behavioral data and custom MATLAB code used for the data analysis of this study will be made available from the corresponding author upon reasonable request.

### Histology

After completion of recording procedures, electrolytic microlesions (12 µA negative current between one wire of each stereotrode and the ground, 30 s duration) were made at the tips of all stereotrodes using the nanoZ (Multi Channel Systems, USA). The next day, the rats were deeply anesthetized with ketamine (75 mg/kg; i.p.) and xylazine (10 mg/kg; i.p.), and then transcardially perfused with 300 ml saline followed by 300 ml 4% paraformaldehyde. The brains were removed and 40 µm slices were processed for cresyl violet staining. The final locations of the electrodes were verified.

## Results

### Behavioral task performance

Five rats were trained in one of two versions of the 8-arm radial maze task, the same-reward (n=2) and the different-reward version (n=3) (Fig. 1A). After extensive training (40 − 50 days), the rats rarely visited the non-reward arms. Furthermore, after consuming the reward in one arm, rats only occasionally revisited the arm in the same trial. Therefore, for each trial only the three approach runs towards the end of the reward arms were considered, while the runs to non-reward arms were not analyzed. Because travelled distances from the starting location to the reward location were long and constant, speed varied from locomotion start to locomotion end in a very stereotyped way (Fig. 1B). Each run consisted of a single acceleration to peak speed followed by a deceleration to locomotion end at the reward location (i.e., along the way to the end of the run, rats did not stop and restart, or slow down and re-accelerate). This consistent pattern of speed variation during each run allowed us to evaluate the relationship between kinematic variables and neuronal activity. Unlike most tasks used to study approach behavior, no start cue was provided and each run was therefore self-initiated.

The two versions of the 8-arm radial maze used in this study were chosen to test whether differences in expected value and arm preference were encoded by NAc neurons. We used the number of times each arm was visited first in each trial as a preference score. Fisher test was used to compare the number of trials each rat first entered each arm versus the number of times they were expected to enter if they had no preference (i.e., 1/3 of the trials). One of the two rats trained under the same-reward version entered significantly more times in arm X (P < 0.01) as first choice but did not show preference for entering arm Y or Z as first choice (Fig. 1C). The other rat trained under the same-reward version did not show significant preference for entering any of the reward arms (P > 0.15). The three rats trained under the different-reward version showed on average a significant preference (vs. chance) for entering arm M (medium reward arm, baited with four drops of chocolate milk in 2/3 of the trials and no reward in ⅓ of trials, P < 0.001), significantly less preference for entering arm L (low reward arm, baited with one drop of chocolate milk in all trials, P < 0.001), and no preference for entering arm H (high reward arm, baited with four drops of chocolate milk in all trials, P = 0.14). This suggests that a reward of low magnitude decreased the rats’ preference, but their preference was not affected by a lower reward probability. The rats’ overall preference for entering arm M first rather than arm H may be explained by the pre-training procedure, in which the arm that was least preferred when the three arms were baited with the same reward was assigned as arm H and the most preferred arm was assigned as arm L in the subsequent training stage.

### Speed and fraction of the run are the best predictors of NAc firing rate during approach behavior

Sixty-two NAc neurons were recorded while trained rats performed the radial arm maze tasks (Tab. 2). To examine how NAc neurons encode reward and kinematic parameters of locomotion, we considered the set of variables shown in Table 1. Many of these were calculated from video tracking data and changed from moment to moment within a single run (e.g., speed, distance traveled); others were constant throughout each run but changed across runs (e.g., trial number, maximum speed during the run). We tested whether the firing rate of each neuron could be predicted with a general linear model (GLM) that used all these parameters as predictive variables. Before running the GLMs, we examined the correlation between pairs of variables and found that fraction of the run, time and distance travelled were strongly correlated with each other (r > 0.8, Fig. 2). We therefore eliminated time and distance travelled from the GLM analysis. In addition, fraction of the run and acceleration were correlated, but less strongly (r = 0.6). Therefore, we conducted three separate GLM analyses for each neuron, one including both variables and the other two including either fraction of the run or acceleration. We refer to these three GLMs as full models (FMs).

**Table 1.** Description of individual variables considered for use as predictors in the GLM encoding model. Variable types include Continuous (continuous variables that could change throughout each run), Constant (continuous variables that were constant throughout each run, but could change across runs), and Dummy (0 or 1). Variables marked Yes under In GLM were included in the GLMs; those marked No were not. “Arm Y” and “Arm Z” were both 0 in cases where the animal entered Arm X. For rats in the different-reward group, Arms X, Y and Z refer to Arms L, M and H, respectively.

|  | Variable type | In GLM | Unit or value | Description |
| --- | --- | --- | --- | --- |
| **Speed** | Continuous | Yes | cm/s | Movement speed of the animal during the task |
| **Fraction of the run\*** | Continuous | Yes | from 0 to 1 | Fraction of the total distance traveled during the reward approach |
| **Acceleration** | Continuous | Yes | cm<sup>2</sup>/s | Acceleration of the animal during the task |
| **Max speed** | Constant | Yes | cm/s | Maximum speed reached during the run |
| **Arm Y** | Dummy | Yes | 1 or 0 | 1 if the animal is approaching reward arm Y |
| **Arm Z** | Dummy | Yes | 1 or 0 | 1 if the animal is approaching reward arm Z |
| **Run duration** | Constant | Yes | s | Total duration of the run |
| **Trial** | Constant | Yes | from 1 - 12 | Number of the trial in a session of 12 trials |
| **Run** | Constant | No | from 1 - 36 | Number of the run in a session of 12 trials |
| **Angle - center** | Continuous | Yes | degrees (0 - 360) | Angle formed by the center of the maze and the animals location |
| **Head direction** | Continuous | Yes | degrees (0 - 360) | Animal's head direction |
| **Head movement** | Continuous | Yes | degrees (-360 to +360) | Changes in head direction (counter-clockwise or clockwise) |
| **Distance traveled\*** | Continuous | No | cm | Distance traveled since the locomotion onset of each run |
| **Time\*** | Continuous | No | s | Time since the locomotion onset of each run |
| **Center distance** | Continuous | Yes | cm | Distance between the animal location and the center of the maze |

\*The values of these variables are reseted (to zero) at the beginning of each new run

We first assessed the validity of the GLM that included both fraction of the run and acceleration using a five-fold validation method (see Engelhard et al. 2019). For each neuron, we computed the r^2^_mean_: the mean of the five r^2^ values from the correlations between the modeled firing rates (obtained from running the GLM on ⅘ ofthe data) and the rem aining ⅕ of the actual firing rate data (Fig. 3). Representative examples comparing actual firing rate with the firing rate predicted by the FM are shown in Fig. 4B. The average r^2^ across the 39 neurons (r^2^_mean_ ± SEM) was 0.15 ± 0.014. We then ran the model 1000 times for each neuron on shuffled data sets and obtained the r^2^_mean_ value from each shuffled data run. The number of neurons per rat with variance explained by the FMs is shown in Table 2. FMs predicted the changes in the firing rate of 39 units (∼63% of the neurons) with a r^2^_mean_ greater than the correlation predicted by 99% of the models using shuffled data (Fig. 4A). In the model in which the variable acceleration was excluded, 39 cells (63%) passed this criterion (Fig. 4-1) and in the model in which the variable fraction of the run was excluded 36 cells (58%) passed this criterion (Fig. 4-2). Therefore, in the majority of neurons, our GLMs predicted firing rates much better than chance, and in these neurons the models were able to account for approximately 15% of the variance in firing rate.

**Table 2:** Summary of recorded cells in individual rats.

|  | Rat #04 | Rat #08 | Rat #09 | Rat #11 | Rat #19 | Total | % |
| --- | --- | --- | --- | --- | --- | --- | --- |
| Number of recorded neurons | 18 | 4 | 25 | 11 | 4 | 62 | 100 |
| Number of neurons with signifcant GLM result | 16 | 1 | 12 | 9 | 1 | 39 | 63 |
| Number of locomotion-off cells | 3 | 2 | 7 | 1 | 1 | 14 | 23 |
| Number of peak-valley cells | 10 | 0 | 4 | 1 | 0 | 15 | 24 |
| Number of valley-peak cells | 2 | 1 | 4 | 5 | 0 | 12 | 19 |
| Number of unclassified cells | 3 | 1 | 10 | 4 | 3 | 21 | 34 |
| Total | 18 | 4 | 25 | 11 | 4 | 62 | 100 |

**Figure 4:**
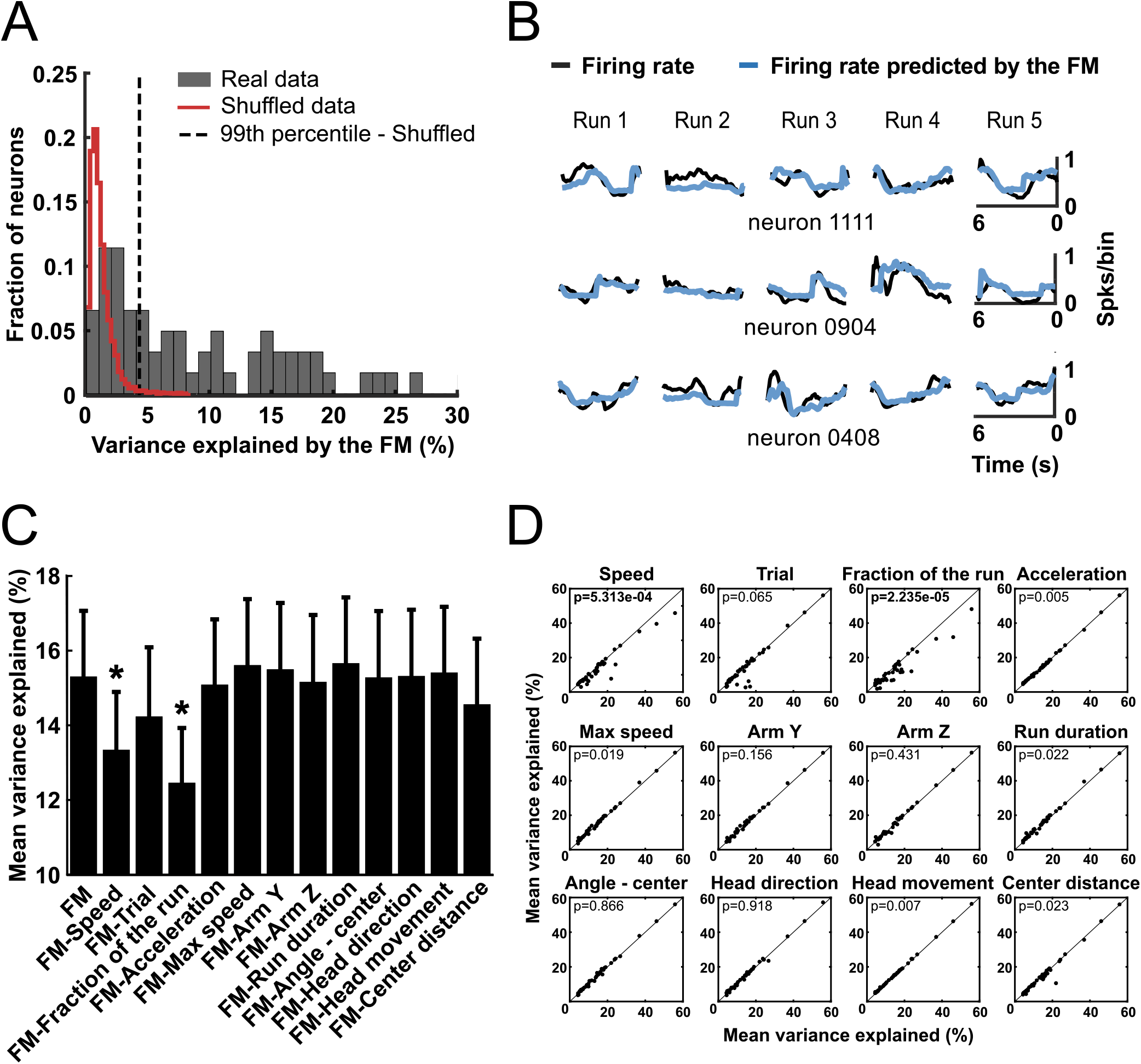
Generalized linear models (GLM) were used to predict activity of NAc neurons based on behavioral variables. **(A)** Distribution of the variance explained by a GLM full model (FM) that used all variables shown in (C) as predictors. Red line delimits the distribution of the variance explained by a GLM calculated with shuffled neuronal data (100 shuffled models per neuron). Neurons on the right of the dashed line (99th percentile of the shuffled models distribution) were classified as significantly explained by the GLM (63% of recorded neurons). In this and subsequent graphs, variance explained is depicted as a percent (100 x r^2^_mean_). **(B)** Neuronal activity during 5 runs towards the ends of reward arms predicted by the GLM (blue line) compared to the real data (black line) for three individual neurons. **(C)** Percentage of mean variance explained (mean ± SEM of r^2^_mean_ across neurons). Only data from the neurons with activity significantly predicted by the FM (full GLM model) were included. The mean variance explained by the FM is shown in the first column. The other columns show the activity explained by the FM models excluding the indicated variable. * p < 0.01, paired t-test (after Bonferroni correction) comparing the indicated model with the FM. **(D)** Comparison of mean variance explained by the FM (X-axis) and by the FM after excluding the indicated variable (Y-axis) for all neurons (black dots) explained significantly by the FM.

To determine which independent variables accounted for the most variance in firing rate, for each neuron we ran the GLM with one variable excluded, and repeated this with each variable. Removing variables that account for the most variance in firing rate should result in the largest reduction in total variance explained. Removing only two individual variables, speed and fraction of the run, significantly decreased the r^2^_mean_ across neurons (P < 0.01, t-test, Fig. 4C,D). Removing acceleration had an effect that did not reach significance after correction for multiple comparisons (Fig. 4D); however, when the variable fraction of the run (which was correlated with acceleration, r= 0.6) was excluded from the model, removing acceleration caused a significant decrease in r^2^_mean_ (Fig. 4-2). Furthermore, when acceleration was excluded from the model, removing the variable fraction of the run caused an even higher decrease in r^2^_mean_ (Fig. 4-1). This suggests that part of the predictive information carried by the variable fraction of the run that is encoded by the NAc neurons is the change in acceleration during the run.

Finally, to confirm that speed and fraction of the run strongly influence firing, we ran GLMs that included only these two independent variables. These models predicted the firing rate of 38 (61%) of the neurons with a r^2^_mean_ greater than the coefficient generated in 99% of models using shuffled data. Thus, our GLM analysis reveals that at least two kinematic variables (speed as well as the progression of the animal along the run and/or acceleration) are most strongly predictive of NAc firing activity, whereas other variables (e.g., those related to the animal’s choice of arm to enter) are not consistently related to firing activity of the NAc neurons.

Although our GLM analysis showed that the arm variable (the arm the animal entered) was not necessary to predict the firing rate of most recorded neurons, we further tested whether arm choice affected neuronal activity in different phases of the approach behavior. First, for each neuron, we pooled data from all runs towards the same arm. The mean firing rate was calculated during four phases: the pre-locomotion phase (the 3 seconds before locomotion start), the acceleration phase (from locomotion start to the speed peak time), the deceleration phase (from the peak speed time to locomotion end), and the post-locomotion phase (the 3 seconds after locomotion ended). Runs with inter-trial intervals shorter than 6 s were excluded. We then compared firing rate in these phases with two-way ANOVA taking the phases as repeated measures and the target arm as the other independent variable (Fig. 6-2). The phase factor was significantly different for 44 out of 62 neurons . The arm factor was significantly different for only three neurons, and the interaction between these factors was significant for eight neurons (P < 0.01). The fraction of neurons that showed significant arm or interaction effects in rats trained under different rewards in the three reward arms was not significantly different from rats trained under the same-reward condition (Fisher test, P < 0.01). This finding reinforces the hypothesis that most of the recorded neurons did not encode the arm choice or reward expectation. Based on this finding and also because the GLM analysis indicated that encoding of the arm entered was weak or non-existent (Fig. 4), in the subsequent analyses data from all runs were pooled. This approach is further justified by our observation (described above) that arm H was not strongly preferred by the group of rats receiving different rewards in different arms.

### Three distinct activity patterns

A neuron whose firing is related to the variable fraction of the run could encode a number of biologically-relevant parameters that themselves vary according to the fraction of the run completed, such as reward proximity and acceleration. Another possibility is that the fraction of the run neurons reach a peak at a specific relative location along the approach trajectory, which could occur at locomotion onset, locomotion offset, or any point in between. To explore these possibilities we first constructed z-normalized histograms of each neuron’s activity aligned to locomotion onset, peak of acceleration, peak of speed, peak of deceleration and locomotion end. These histograms are plotted as heat maps sorted by the time at which the peak activity occurred (Fig. 5A). Some neurons seemed to be inhibited during the runs, an observation we confirmed with paired t-tests comparing the firing rate during the runs with the firing rate during the inter-run intervals. Fourteen neurons were significantly inhibited during the locomotion phase (P < 0.01) and were named **Locomotion-Off (LO) cells**. An example neuron’s raster and histogram are shown in Fig. 6A1,2, and histograms of all LO cells are shown in Fig. 7A. These histograms show that LO cells tended to be inhibited beginning just prior to locomotion onset, continue to be inhibited throughout the run, and abruptly recover from the inhibition just after locomotion offset.

**Figure 5:**
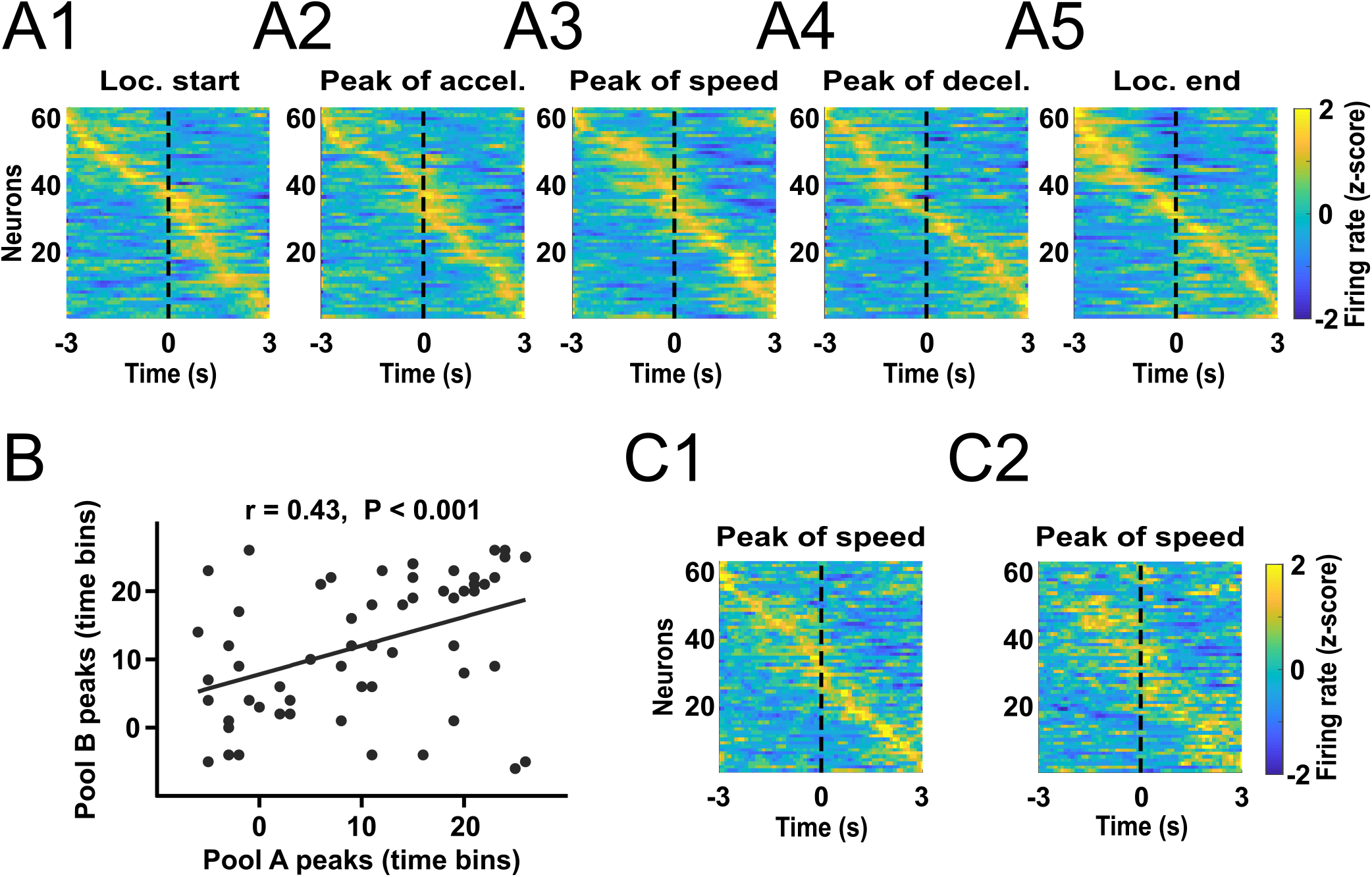
Different neuron peak activity at different bins of time. **(A)** Averaged firing rate z-score histograms from all runs of all recorded neurons aligned to locomotion start (A1), peak of acceleration (A2), peak of speed (A3), peak of deceleration (A4), locomotion end (A5). **(B, C)** Firing rate z-score data from all runs were aligned to the peak of speed and split into two pools counterbalanced for the trial order and visited arm. The time bin in which the activity of each neuron peaked was calculated separately for the two pools and plotted against each other. Statistical results are from the Pearson test for correlation. **(C)** Neurons were sorted by the peak of activity calculated with the first data pool (C1). The activities of the same neurons calculated with data from the second data pool is shown in the same order as for the first data pool (C2).

**Figure 6:**
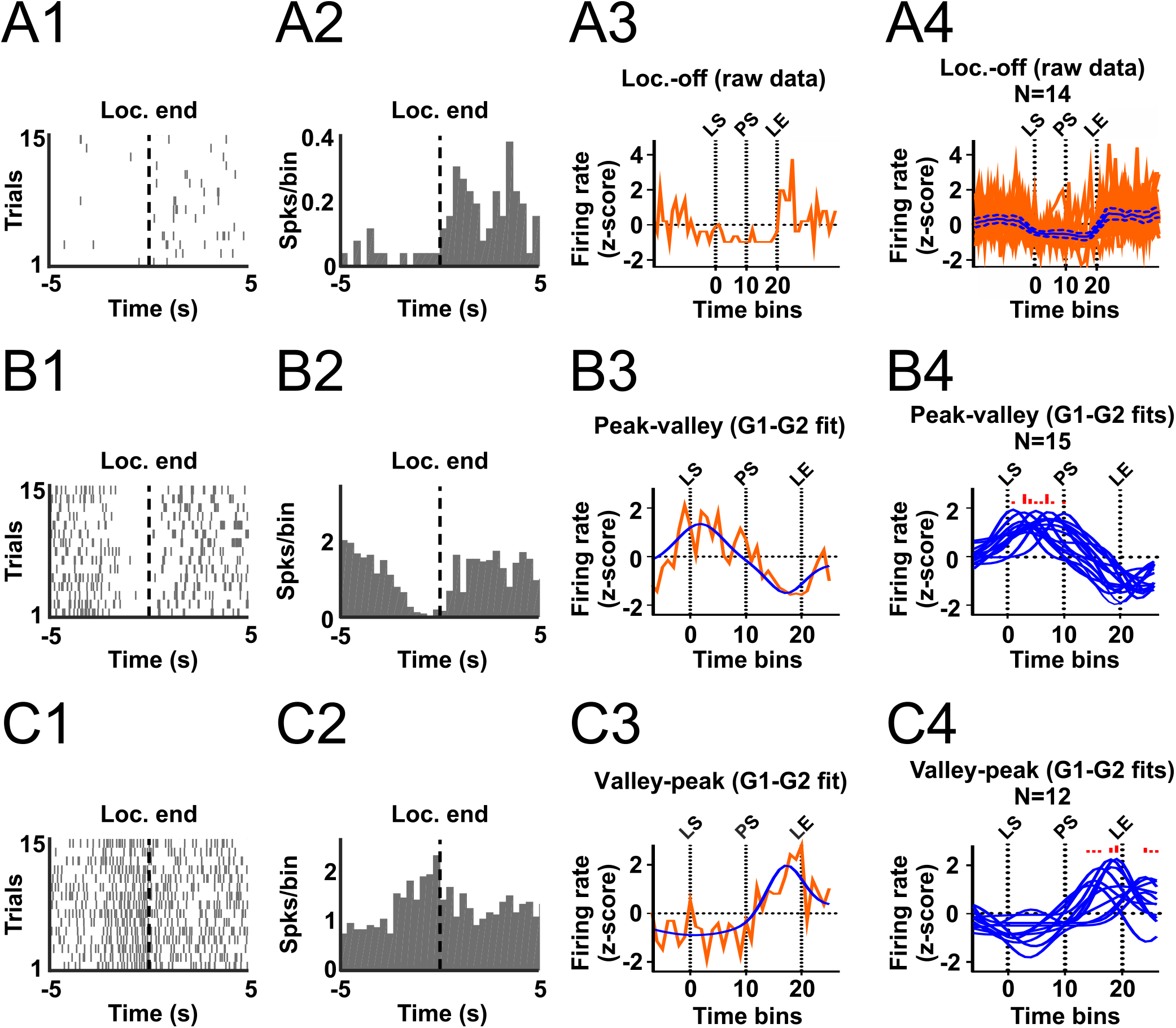
Classification of LO, PV and VP cells. **A1** and **A2** show rasters and histograms aligned to locomotion end for an example LO cell. **A3** shows the time-normalized histogram for this neuron, and **A4** shows time-normalized histograms for all LO cells superimposed (red traces) and the mean ± SEM (blue trace and blue dashed traces). **B** and **C** show similar plots for PV (**B**) and VP (**C**) neurons. The blue traces in **B3, B4, C3** and **C4** show individual neurons’ 2G model results. The red histograms superimposed above **B4** and **C4** indicate the time at which each neuron’s peak firing occurs according to its 2G model. LS, PS, and LE correspond to locomotion start, peak of speed, and locomotion end, respectively.

**Figure 7:**
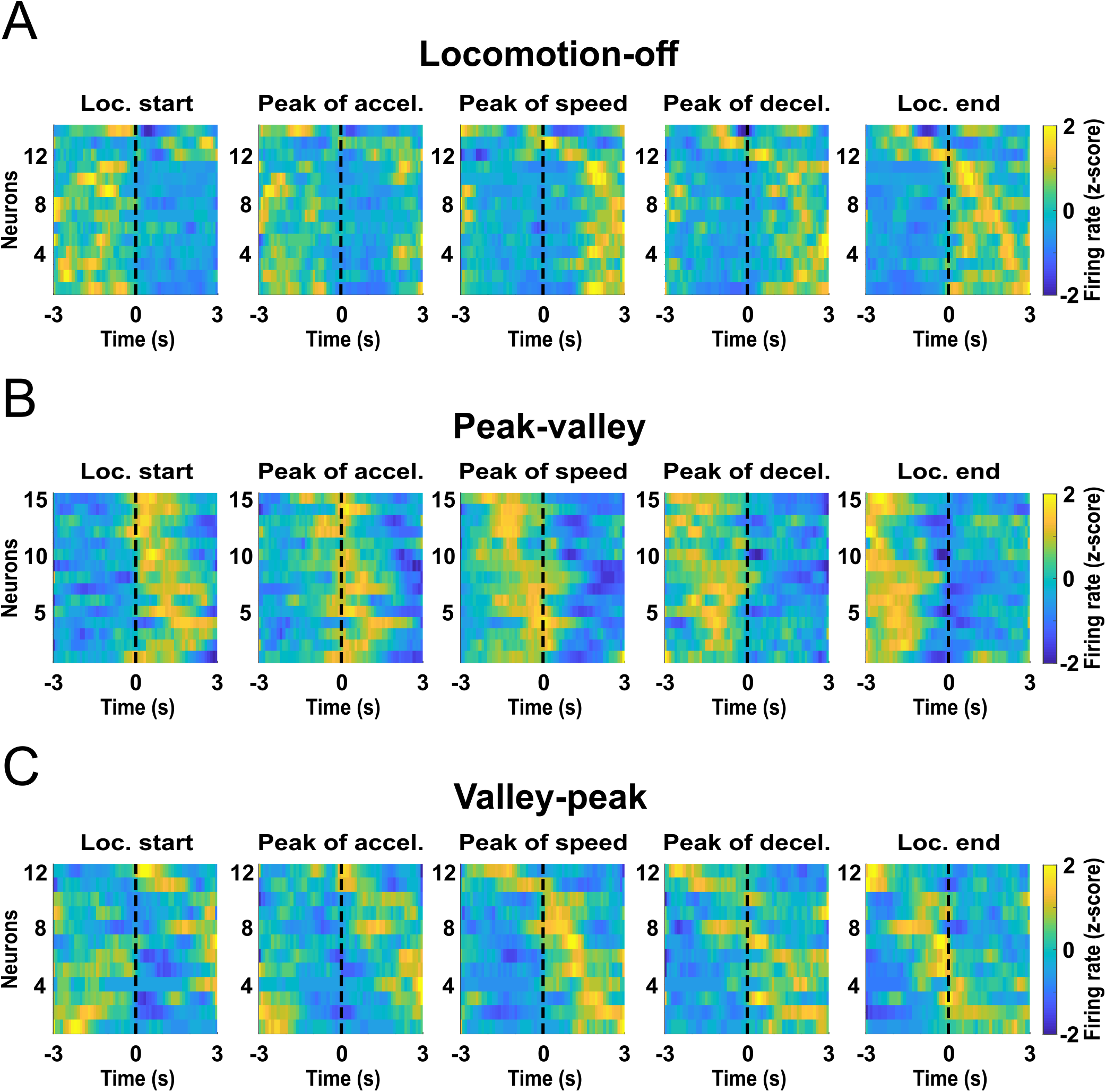
LO (A), PV (B) and VP (C) cells exhibit peak activity at different kinematic stages of the reward approach run. Histograms show firing rates aligned to locomotion start, peak of acceleration, peak of speed, peak of deceleration, and locomotion end. Data were z-scored and the colors in each row show the average across runs for an individual neuron.

Fig. 5A shows that many of the remaining neurons exhibited activity peaks at various times from before locomotion start to after locomotion end. Consequently, the distribution of the histogram peaks sorted by time from the alignment event formed a diagonal line, particularly when activity was centered on the peak of speed (roughly the midpoint of the run). To assess whether this diagonal was the result of each neuron’s firing consistently reaching a peak at about the same time across runs relative to the peak of speed, we split the data from each neuron into two pools counterbalanced by trial order and target arm. Peak speed-aligned histograms of the activity of all neurons calculated with the first data pool, sorted by the time of peak of activity, are shown in Fig. 5C1; and the histograms calculated with the second data pool, sorted in the same order as the first pool, are shown in the Fig. 5C2. We found that the times of the peak bins determined from the two data pools were correlated (Fig. 5B, r = 0.43, p < 0.0001). We also found that for 37 (60%) neurons, the firing rates in each time bin within individual runs were also correlated between data pools (P < 0.01). These results demonstrate that each individual neuron exhibits peaks at consistent times, and that the times during the run at which these peaks occur vary widely across neurons.

Intriguingly, neurons that showed an activity peak in the first part of the approach run (before reaching peak speed; i.e., during acceleration) tended to show a valley in the second part of the run (after peak speed; i.e., during deceleration), and vice-versa (Fig. 5A). We hypothesized that a large subset of neurons exhibited the **Peak-Valley (PV)** pattern of activity because they reach activity peaks and valleys exclusively during acceleration and deceleration, respectively. Similarly, we hypothesized that neurons exhibiting the complementary **Valley-Peak (VP)** pattern reached activity peaks and valleys exclusively during deceleration and acceleration, respectively. To test these hypotheses, we first had to account for the fact that the durations of the acceleration and deceleration phases were not identical across runs, which means that if firing peaks occurred at a specific fraction of the acceleration or deceleration phase, the peaks would occur at different absolute time points in different runs. Therefore, we time-normalized the firing rate data. In each run, the acceleration period was divided into 10 bins of the same size, and the firing rate in each bin was calculated. The deceleration period was also divided into 10 bins. The approximate average width of the adjusted bins was 150 ms. In addition, 7 bins of 150 ms each were included prior to locomotor start and after locomotor end. Next, the firing rates from all runs were averaged per bin and z-transformed. Finally, we modeled the peak-valley and valley-peak patterns as the sum of two Gaussian curves (2G model, see Methods and Fig. 6-1 for details).

To validate the 2G model, we split the firing rate data from all runs of each neuron into two pools. The first data pool was used to calculate the average activity per bin and the second data pool was used to train the 2G model. Next, we examined whether the activity per bin calculated with the first data pool was significantly correlated with the activity predicted by the 2G model (Pearson correlation, P < 0.01). Fifteen PV cells (24%) and 12 VP cells (19%) passed this criterion (Fig. 6B,C). Note that the peak firing rate of PV cells occurred exclusively during the acceleration phase (Fig. 6B4, 7B), whereas the firing rate peak of VP cells occurred mostly during the deceleration phase, although some neurons peaked after locomotion end (Fig. 6C4, 7C). LO cells were not modelled with the 2G model, but we plotted the time-normalized firing rates of these neurons in the same way as for the PV and VP cells in Fig. 6A4.

These firing modes further refine the GLM results (Fig. 4C,D) showing that speed and fraction of the run (or acceleration) explain a portion of the variance in firing rate. To examine how LO, PV and VP cells encode speed and acceleration, we plotted the simple correlation coefficients relating firing rate and speed against the normalized times at which the peak firing of each neuron occurred (Fig. 8A), and similarly for the firing rate vs acceleration correlation coefficients (Fig. 8B). PV cells tended to have the strongest positive correlation coefficients for both speed and acceleration, whereas LO neurons had the strongest negative coefficients for speed. VP cells tended to have the strongest negative correlation coefficients for acceleration whereas their coefficients for speed were more widely distributed across negative and positive values.

**Figure 8:**
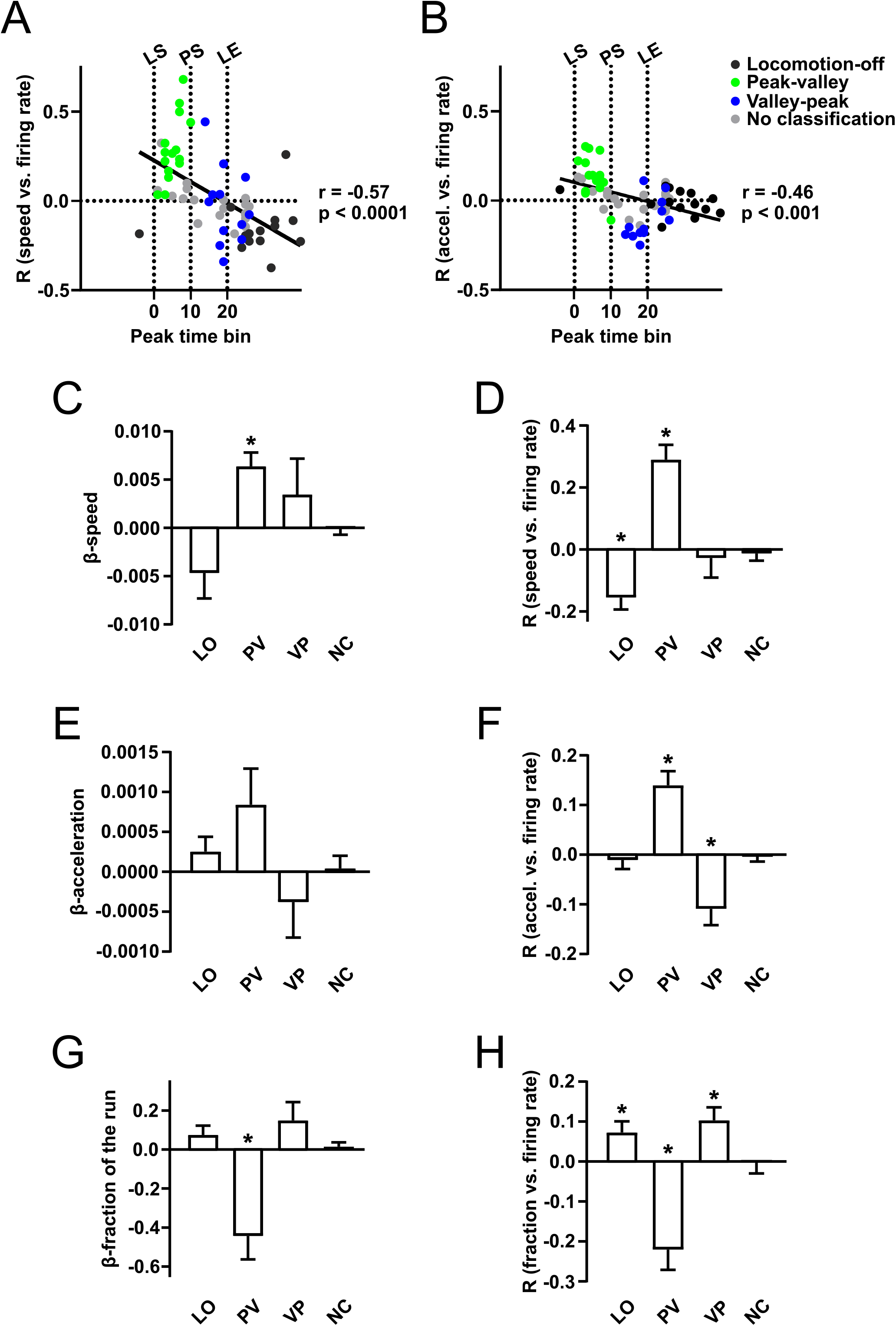
Neurons in each class defined by the 2G analysis share common correlates to predictors in the GLM analysis. The r values for the correlation between speed and firing rate (**A**) and for acceleration and firing rate (**B**) are plotted against the normalized time at which the neuron exhibits peak firing activity. The time of peak is negatively correlated with both r values; the indicated P values are from Pearson’s correlations. Points are color-coded according to classification from the 2G analysis. In **C, E** and **G**, the mean ± SEM weights (*β* values) from the GLM for the variables speed (C), acceleration (E), and fraction of the run (G) are shown separately for LO, PV, VP and non-categorized neurons. Similarly, in **D, F** and **H**, mean ± SEM r values for simple correlations between these variables and firing rate are shown. * P < 0.01, one-sample t-test comparison to 0.

To further confirm these findings based on individual neuronal data, we determined, for each class of neurons (LO, PV, VP, and non-categorized), the mean GLM *β* values for speed and acceleration (Fig. 8C,E), as well as the mean r values for simple correlations between speed or acceleration and firing rate (Fig. 8D,F). The results show that PV cells encode speed most strongly, with mean positive coefficients that were significantly different from 0 (P < 0.01, one-sample t test, Fig. 8C,D). In contrast, VP cells did not exhibit consistently negative or positive coefficients for speed, whereas LO cells tended to exhibit negative correlations (Fig. 8C,D). On average, PV cells had significantly positive firing vs acceleration correlations, whereas VP cells’ acceleration correlations were significantly negative (Fig. 8F). These results were recapitulated in the GLM *β* values for acceleration, although they did not reach significance (Fig. 8E). On average, the correlation coefficients for fraction of the run were also significant (and the GLM *β* values tended to significance) for the 3 classes of cells (Fig. 8G,H). For PV and VP cells, the signs of the fraction of the run coefficients were the opposite of those for acceleration, as expected given that acceleration occurs early in the run and deceleration occurs later.

Together, these results indicate that the encoding of acceleration and/or fraction of the run revealed by the GLM is largely the result of PV and VP cells’ firing peaks and valleys, whereas encoding of speed is largely due to PV cells’ positive correlations of firing with speed and LO cells’ with negative correlations.

### Activity of speed-correlated neurons precedes increases in speed

Our finding that the activity of NAc neurons is correlated with approach speed is in agreement with the hypothesis that NAc neurons promote vigorous reward seeking (Nicola, 2007; Salamone and Correa, 2012; Nicola, 2016). If the firing of these neurons is causal to increased speed of reward approach, then increases in speed should reliably follow the activation of these neurons. We tested this prediction with a cross-correlation analysis in which we plotted speed versus time aligned to each spike. Neurons were sorted according to the correlation between speed and firing rate (Fig. 9A1). On average, the speed of the positively-correlated neurons peaked 146 ms after the action potential (P = 0.0017, t test for difference from 0, Fig. 9B). In contrast, the peak of the cross-correlograms of the negatively-correlated neurons were not significantly different from the spike times (P = 0.35, Fig. 9B). As suggested by Fig. 8, PV cells were the class with the strongest correlation between firing and speed. To verify that the firing of these neurons preceded an increase in speed, we constructed heat maps of LO, PV and LO cells’ firing-speed cross-correlations. These plots show that only PV cells consistently exhibited cross-correlogram speed peaks after the spike (Fig. 9A2). Notably, the cross-correlogram speed minimum tended to follow the spike in VP cells. These results are consistent with the possibility that the firing of PV cells causes an increase in the speed during the acceleration phases, whereas the firing of some VP cells causes a decrease in speed during the deceleration phase.

**Figure 9:**
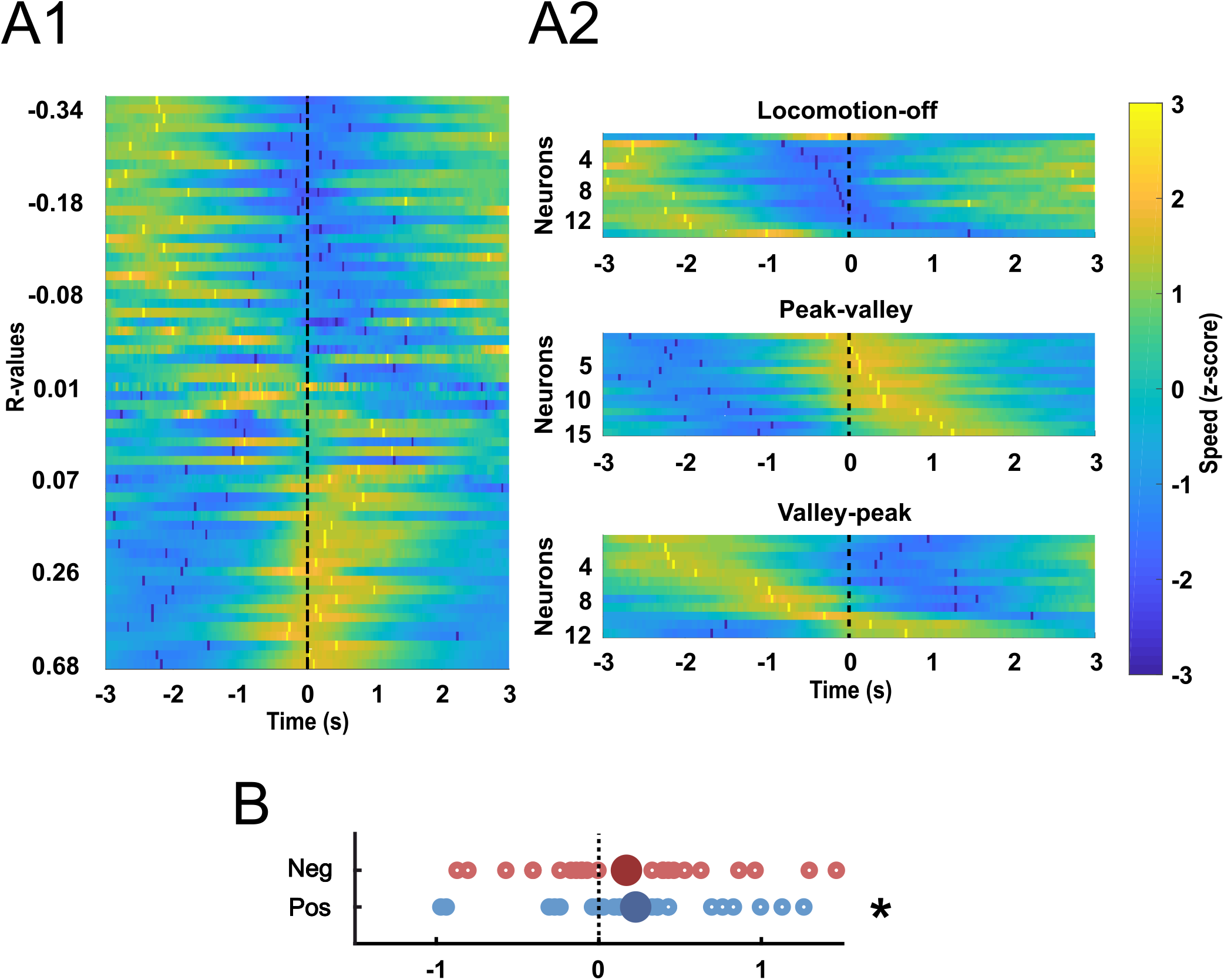
Spike-speed cross-correlations. **A1**, Each row contains a histogram of the average z-score (colors) of speed aligned to the spikes of an individual neuron. The bright yellow and dark blue markers indicate peaks and valleys of the cross-correlogram, respectively. Neurons in the heat map were sorted according to the Pearson’s correlation coefficient (r) for speed vs firing rate. All 62 neurons are shown. **A2**, Cross-correlograms are shown separately for LO, PV and LO cells, sorted according to the time at which the cross-correlogram peak speed occurred. **B**, The time at which the lowest cross-correlogram speed occurred is shown for neurons that had negative correlations with speed (top, red points), and the time at which the highest cross-correlogram speed occurred is shown for neurons that had positive correlations with speed (bottom, blue points). Only data from the neurons with significant positive (blue) or negative (red) correlations with speed are shown. * P = 0.04, one-sample t test compared to 0.

### Place-related activity

Alterations in the firing rate of NAc neurons prior to arrival at a reward place have been previously reported (Martin and Ono, 2000; Miyazaki et al., 1998). To test whether the speed or run fraction encoding we identified could be accounted for by location or reward prediction encoding, we performed several analyses. First, we included variables related to location in the GLM described in Fig. 2 including head direction, head movement, angle of the arm and distance from the center of the arm. In the population of neurons as a whole, none of these variables were found to contribute significantly to the model’s predictive power (Fig. 4C). Second, we asked whether any of the recorded neurons could be classified as place cells according to the criterion used by Yeshenko et al. (2004). Five PV and two LO neurons passed the criterion. However, their tentative firing fields were mainly located on the central platform, the neurons were quite active on other parts of the maze, and their firing fields were quite small compared to those of CA1 place cells recorded on a similar radial eight-arm maze (Xu et al., 2019).

### Neuronal classification based on electrophysiological properties

We sorted the recorded neurons into tentative populations of medium spiny neurons (MSNs) and interneurons based on average firing rate and peak width, similar to previous studies (e.g. Berke et al., 2004; Yamin et al., 2013). We classified 34 neurons (55%) as putative MSNs, 8 neurons (13%) as putative interneurons, and 20 neurons (32%) as cells with no clear classification (Fig. 6-3A). 9/14 LO, 3/15 PV, and 8/12 VP cells were classified as MSNs. 1/14 LO, 3/15 PV, and 2/12 VP cells were classified as interneurons. 4/14 LO, 9/15 PV, and 2/12 VP cells showed no clear classification (Fig. 6-3B).

### Histology

Most of the recorded neurons were in the NAc shell, but a few of them were in the NAc core or in the border between the dorsal and ventral striatum (Fig. 6-4). We found no evidence that the different classes of neurons were anatomically clustered.

## Discussion

The present study clarifies how NAc neurons represent the vigor of spontaneous approaches to rewarded locations. Rats had to run long distances (∼1.5 m) to approach each reward, which allowed us to determine how firing changes during large increases and decreases in approach speed. Firing reflected the animal’s speed as well as its progression towards the movement target but not location of the movement target or predicted reward value. Our results suggest that NAc neurons govern the kinematics of approach behavior - when to speed up and when to slow down, and by how much - but, in our task, NAc neurons do not contribute to deciding which target to approach.

We included a broad range of variables related to approach vigor, navigation, and timing in each neuron’s GLM. Only three - speed, fraction of the run completed and acceleration - were found to contribute significantly to the variance in firing rate of the population of NAc neurons (although because acceleration and fraction of the run were correlated, acceleration was found to contribute to variance only when fraction of the run was excluded from the GLMs). Because these variables are related to vigor and kinematics, we conclude that the primary function of NAc neurons in this task is to control the time course of increases and decreases in speed rather than the direction of locomotion.

NAc neurons did not encode kinematic parameters in a uniform way, but rather fell into three classes defined by the trajectory of their firing rates during runs. First, we noted that correlation coefficients relating firing and speed were widely distributed across both positive and negative values. Many neurons with negative coefficients were found to be continuously inhibited during locomotion (LO cells, Figs. 6A, 7A, 8C,D), similar to previously-identified NAc neurons that are inhibited throughout appetitive and consummatory behaviors (Taha and Fields, 2006). However, in LO cells, firing resumed just after locomotion offset even though animals presumably engaged in reward consumption at that point. As suggested previously (Taha and Fields, 2006), the inhibitions of LO cells may gate appetitive behaviors or trigger locomotor approach.

In contrast, many neurons with positive speed coefficients were not continuously excited during locomotion, but rather exhibited firing peaks during acceleration and valleys during deceleration (PV cells, Figs. 6B, 7B, 8A-F). These neurons also tended to exhibit positive correlations with speed/acceleration and negative correlations with fraction of the run completed (Fig. 8E-H). Intriguingly, their firing peaks were distributed throughout the acceleration phase (Figs. 6B4, 8A,B) and did not align precisely to the peak of acceleration (Fig. 7B). These results suggest that PV cells each report that the animal is at a different point along the acceleration trajectory, information that could be used to control the precise timing of the speed increase during the run. This possibility is supported by our cross-correlation results showing that an increase in speed followed a spike in all but one PV cell (Fig. 9A2). This temporal relationship held for most neurons with positive correlations between firing and speed (Fig. 9A1), strengthening the hypothesis that a major role of NAc neurons is to increase the vigor of approach.

The third category of neurons, VP cells, were complementary to PV cells in that VP cells exhibited a firing valley during acceleration. However, VP cells’ peaks were not limited to the deceleration phase, but tended to occur just before or sometimes after locomotion offset (Figs. 6C4, 7B, 8AB). VP cells did not consistently encode speed (Fig. 8A,D), but did tend to show negative correlations with acceleration and positive correlations with fraction of the run completed (Fig. 8E-H) - a pattern opposite to that of PV cells. Thus, PV and VP cells together account for much of the fraction of the run and acceleration encoding identified by the GLM analysis, suggesting that together these neurons report the animal’s relative position along the run. Alternatively, the fact that VP cells tend to fire most near locomotion offset may mean that these neurons are specialized for some aspect of reaching the end of the run, such as coming to a complete stop or preparing for reward consumption.

The GLM analysis yielded little evidence that NAc neurons participate in choosing which reward location to approach or determining the route to get there. In particular, the arm entered variables reflect both movement target and, in some animals, the reward that can be expected, but the firing of few, if any, neurons was influenced by these variables (Fig. 4C,D). Specifically, when the arm variables were removed from the GLMs, the remaining variables explained the variance in firing rate to the same extent as when the arm variables were included. Because this “leave one out” method assesses the contribution of variables to variance independent of whether the variable is positively or negatively correlated, negative results in the cross-neuron analysis are not simply due to existence of roughly equivalent numbers of positively- and negatively-correlated neurons. Rather, variables other than speed, fraction of the run and acceleration did not strongly contribute to the variance of any neurons (Fig. 4D and Extended Figs. 4-1C, 4-2C).

The near absence of representation of navigation and reward parameters contrasts with previous studies showing that NAc neurons integrate reward and spatial information in maze tasks (Floresco et al., 1996; Shibata et al., 2001; Mulder et al., 2004; Mulder et al., 2005; German and Fields, 2007; Ito et al., 2008; Khamassi et al., 2008; van der Meer and Redish, 2009; van der Meer et al., 2010; Lansink et al., 2012) and that the NAc may be required for spatial navigation towards large rewards (Albertin et al., 2000). However, our rats were overtrained and likely approached reward locations (particularly the second and third in each trial) in a sequence of habit-like stimulus-action chains (Graybiel, 1995); i.e., they used a praxic or taxic approach strategy as opposed to a navigational strategy (O’Keefe and Nadel, 1978; Nicola, 2016). Consequently, spatial navigation processing may have been offline or muted, potentially explaining why we found little evidence for location or movement target encoding (McDonald and White, 1993; but see Miyoshi et al., 2012).

Our observation of robust anticipatory speed encoding provides a potential mechanism to explain previous observations indicating a role for NAc neurons in promoting vigorous reward seeking. Approach speed is an important component of vigor (Floresco, 2015; Shadmehr et al., 2016; Salamone et al., 2016), and increasing dopaminergic (Mogenson et al. 1993; Wu et al. 1993) or glutamatergic activity (Maldonado-Irizarry & Kelley 1994) in the NAc increases locomotor activity and invigorates reward seeking (Berridge 2007). Moreover, free-run operant tasks with higher effort requirements are especially susceptible to disruption by interference with NAc dopamine transmission (Salamone et al., 1999). These effects are due to increased latency to return to the operandum after a pause in which the animal moves away from it (Nicola, 2010). One possibility is that dopamine promotes the firing of NAc PV cells, which exhibit the most robust positive correlations between firing rate and speed, and whose firing tends to be followed by an increase in speed. The activity of these neurons could then trigger initiation of locomotor approach by reaching a threshold. PV cells could express the D1 receptor as activation of this receptor tends to have excitatory effects (Andre et al., 2010); because VP cells tend to be inhibited during acceleration, they may express D2 receptors, causing them to be inhibited by dopamine. Recordings from identified D1 and D2 NAc neurons are needed to address these hypotheses.

Consistent with the idea that PV cells drive locomotor initiation, previous studies showed that the magnitude of NAc neuronal excitations in response to reward-predictive cues predicts both the latency to initiate approach locomotion and the speed of approach (Morrison et al., 2017; McGinty et al., 2013). These cue-evoked excitations are brief (typically well under 1 s) and precede initiation of movement. Although this time course contrasts with our observation of speed encoding during locomotion, the previous studies used smaller operant chambers in which locomotor events were brief and higher speeds could not be attained. Thus, it is possible that neurons exhibiting cue-evoked excitations are the same as the PV cells identified here. Alternatively, because cue-evoked excitations are greater when the subject is closer to reward (Morrison 2014, 2017), it is possible that these neurons are VP cells because VP cells tend to exhibit bursts as the animal reaches the rewarded location. Further comparative work is needed to assess whether encoding of kinematic parameters occurs in the same neuronal populations across tasks, and whether similar encoding occurs during locomotion that is not directed towards reward.

Our results demonstrate that NAc neurons’ activity during free-run reward approach is most consistent with control by these neurons over approach kinematics - when to speed up and when to slow down, and by how much. Future studies should use tasks requiring cognitive map-based navigation to assess how encoding of kinematics is related to forms of activity that contribute to deciding where to go and how to get there.

## Acknowledgments

The financial support of the grants CNPq (432061/2018-5, 306855/2017-8, 465346/2014-60), AZV 17-30833A, and GACR 20-00939S, is acknowledged.

