## Supplementary figures and images for "Nucleus accumbens neurons encode initiation and vigor of reward approach behavior"

### Extended Figure 4-1

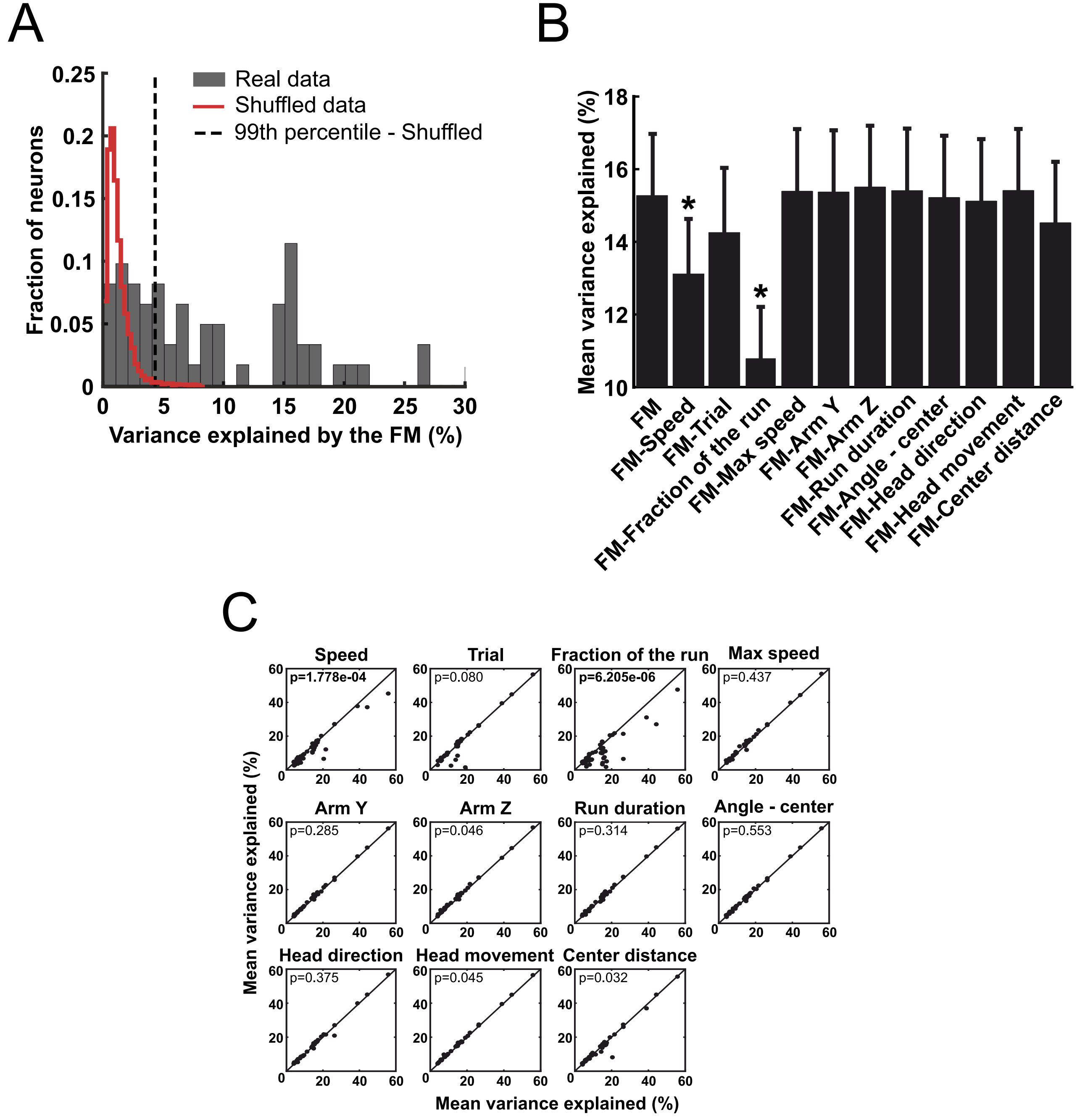

### Extended Figure 4-2

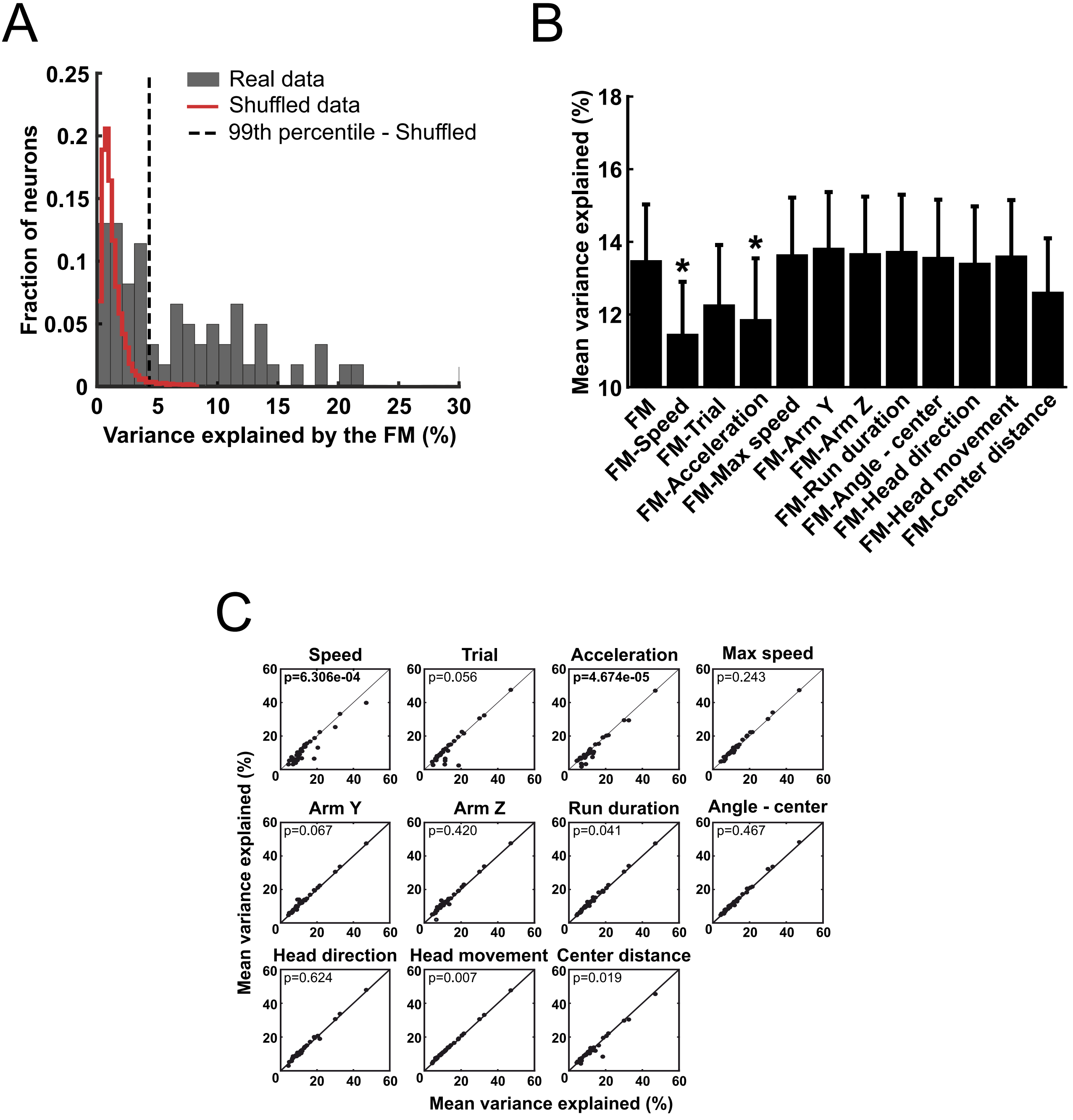

### Extended Figure 6-1

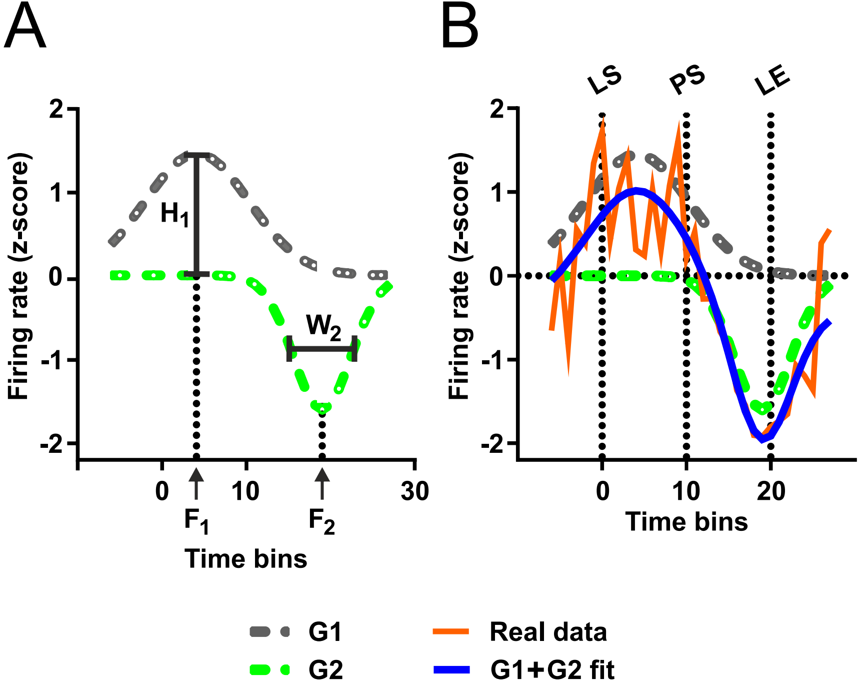

### Extended Figure 6-2

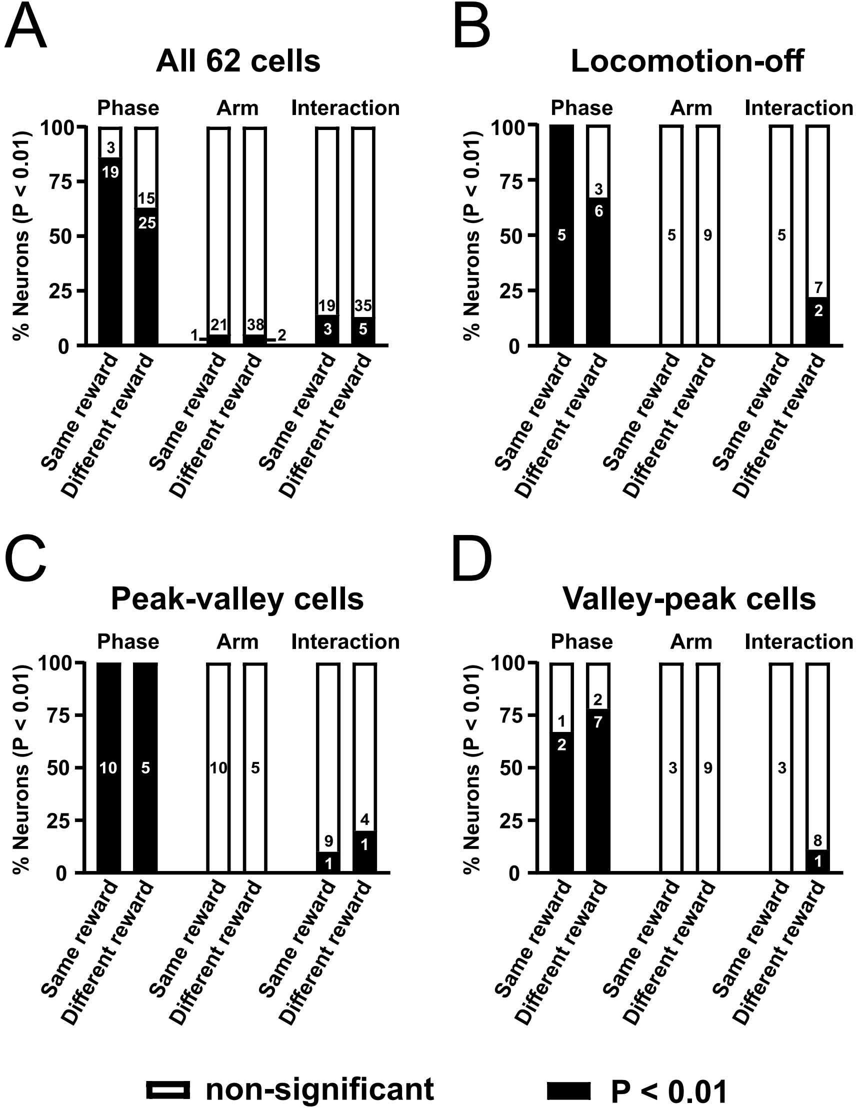

### Extended Figure 6-3

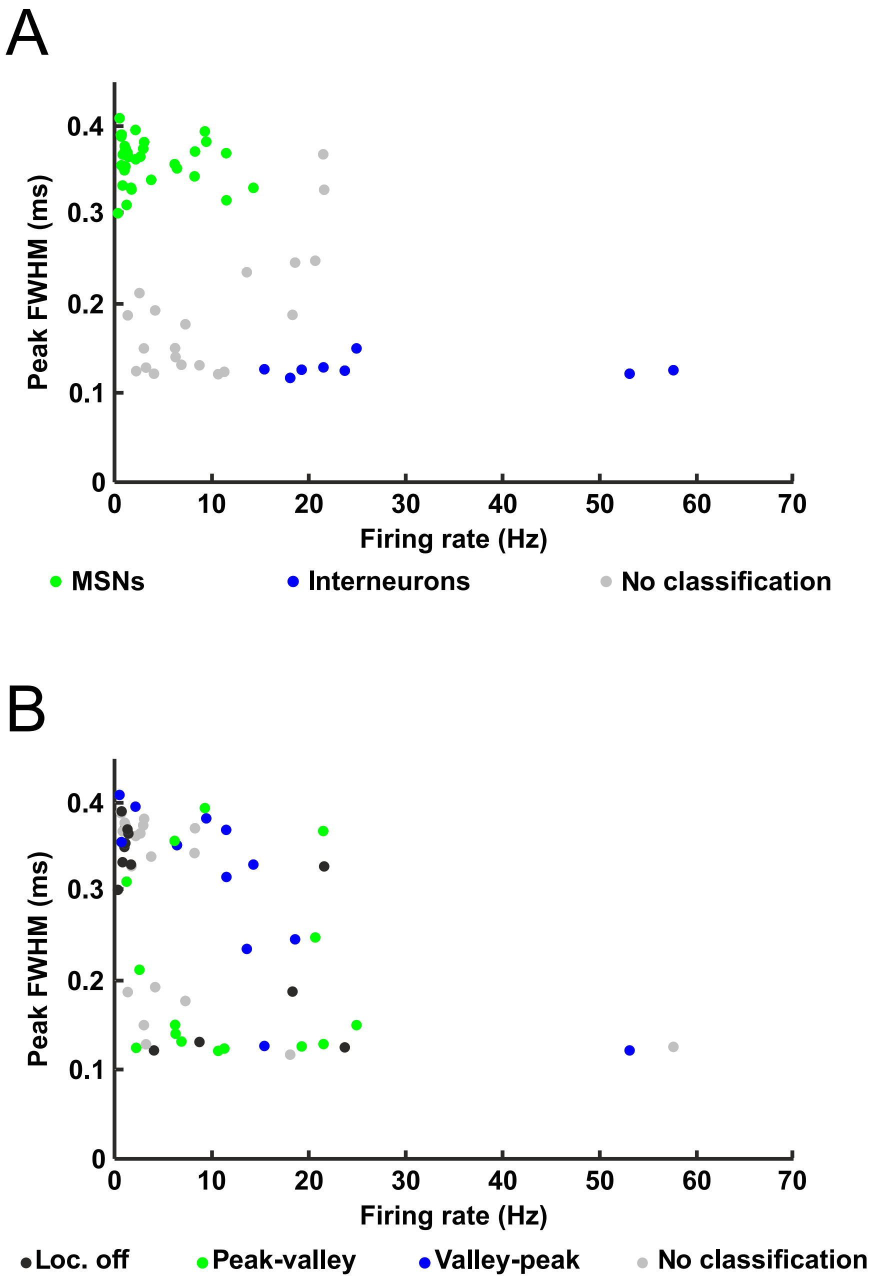

### Extended Figure 6-4

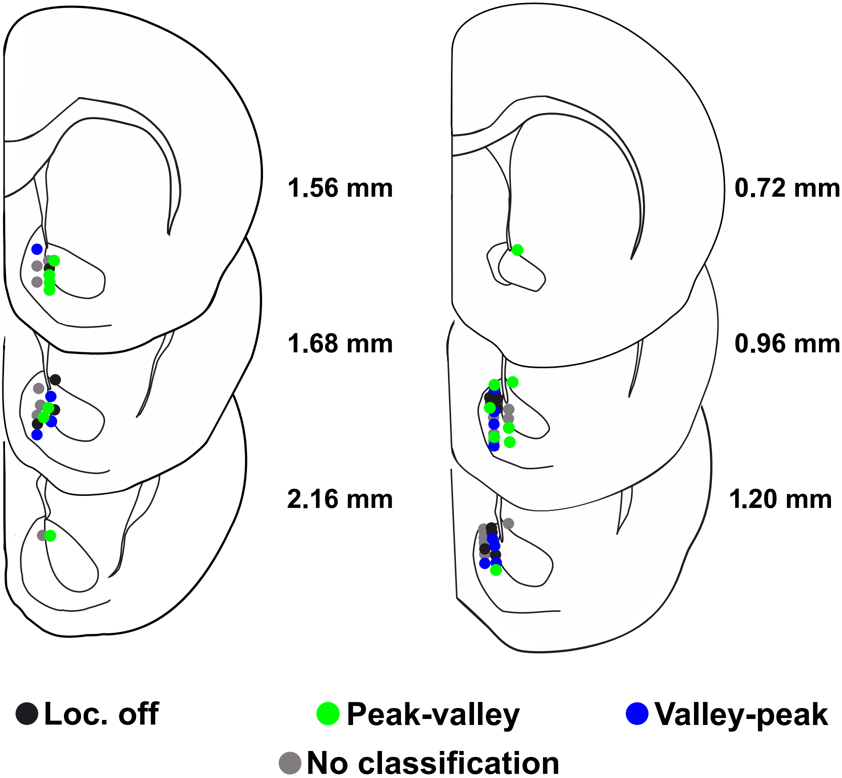
