## Extended Figure legends for "Nucleus accumbens neurons encode initiation and vigor of reward approach behavior"

**Figure 4-1**: Generalized linear models (GLM) without the acceleration variable. **(A)** Distribution of the variance explained by a GLM full model (FM) that used all variables shown in (B) as predictors. Red line delimits the distribution of the variance explained by a GLM calculated with shuffled neuronal data (100 shuffled models per neuron). Neurons to the right of the dashed line (99th percentile of the shuffled models distribution) were classified as significantly explained by the GLM (63% of recorded neurons). In this and subsequent graphs, variance explained is depicted as a percent (100 x $r^2_{mean}$). **(B)** Percentage of mean variance explained (mean ± SEM of $r^2_{mean}$ across neurons) of the firing rate of neurons with activity significantly explained by the FM. The mean variance explained by the FM is shown in the first columns. The other columns show the variance explained by the FM excluding the indicated variable. * $p < 0.01$, paired t-test (after Bonferroni correction) comparing the indicated model to the FM. **(C)** Comparison of mean variance explained by the FM (X-axis) and by the FM excluding the indicated variable (Y-axis) for all neurons (black dots) explained significantly by the FM.

**Figure 4-2**: Generalized linear models (GLM) without the fraction of the run variable. Panels are as described for Fig. 4-1, except that these GLM predicted the changes in the firing rate of 58% of the neurons.

**Figure 6-1**: The sum of two Gaussians (2G) model. A non-linear model that consists of the sum of two Gaussian curves was used to model the firing rate during the approach run. The panel A shows the two modeled Gaussian curves, and the panel B shows the same two curves as well as the mean firing rate data (z-score) and the sum of the two Gaussian curves. The only predictor used in this model is the fraction of time. The period of time from the locomotion start (LS) to the peak of speed (PS) was divided into 10 bins, and the time from the peak of speed and the locomotion end (LE) was also divided into 10 bins. These bins were ~150 ms each. The 7 bins preceding locomotion start (LS) and following locomotion end (LE) were 150 ms each. Changes in z-scored firing rate as function of normalized time ($Y_{(t)}$) was modelled as $Y_{(t)} = G1_{(t)} + G1_{(t)} + BL$, where G1 was the equation for the peak part of the curve, G2 was the equation for the valley part of the curve and BL was the baseline activity. The Gaussian equations ($G_{(t)}$) used were: $G1_{(t)} = H1e^{-\pi(\frac{X_{(t)}-F1}{W1})^2}$, $G2_{(t)} = -H2e^{-\pi(\frac{X_{(t)}-F2}{W2})^2}$, where H, F, and W are constants with the following meaning: H represents the amplitude of the Gaussian curve, which reflects the height (H1) and depth (H2) of the activity peak and the valley, respectively. F is the bin where the Gaussian is centered. Therefore, F1 is the bin in which the neuron is most active, and F2 is the bin in which the neuron is least active. W is the standard deviation of each Gaussian, which reflects the durations of the activation peak (W1) and valley (W2).

**Figure 6-2**: Effects of the target arm and the phases of the approaches run on the firing rate. The firing rates were scored during the 3 s before the approach run and during the acceleration and deceleration phases of the runs. Independent repeated measures (phases) ANOVAS were run for each neuron. Bars represent the percent of neurons with significant phase factor, significant arm factor, and significant interaction phases ($P < 0.01$). The absolute number of neurons in each category is printed inside the bars. No significant difference between the number of neurons with significant effects from rats that were trained with the same and rewards in the 3 arms was found ($P > 0.01$, Fisher test).

**Figure 6-3**: Separation of recorded cells into clusters of tentative MSNs (blue), interneurons (red), and neurons with no classification (gray).

**Figure 6-4**: Placement of tips of recording electrodes. Neurons located outside of the nucleus accumbens were excluded from the analysis.
